# Headache-specific Hyperexcitation Sensitises and Habituates on different Time Scales: An Event Related Potential study of Pattern-Glare

**DOI:** 10.1101/2024.08.28.610154

**Authors:** Cihan Dogan, Claire E. Miller, Tom Jefferis, Margarita Saranti, Austyn J. Tempesta, Andrew J. Schofield, Ramaswamy Palaniappan, Howard Bowman

**Affiliations:** School of Computing, University of Kent, Canterbury, UK; School of Psychology, College of Life and Environmental Sciences, University of Birmingham, Edgbaston, Birmingham, UK; School of Psychology, College of Health and Life Sciences, Aston University, Birmingham, UK; School of Computer Science, University of Birmingham, Edgbaston, Birmingham, UK

## Abstract

Cortical hyperexcitability is a key pathophysiological feature in several neurological disorders, including migraine, epilepsy, tinnitus, and Alzheimer’s disease. We examined the temporal characteristics of Evoked Related Potentials (ERPs) in a healthy population using the Pattern Glare Test, a diagnostic tool used to assess patients with sensitivity to cortical hyperexcitability. During the experiment, participants recorded state measures with this study focussing on susceptibility to migraine. We investigated two timeframes: habituation over the course of the experiment and sensitization over the course of stimulus presentation. We found evidence of hyperexcitability in the visual cortex, for the clinically aggravating stimuli (medium). Participants who reported a higher state measure exhibited a higher degree of habituation and sensitization, which was dependent on susceptibility to migraine. These findings suggest that the same experimental paradigm and analysis should be performed on a clinically diagnosed population.

## 1. Introduction

Some visual stimuli cause mild discomfort for most observers and considerable discomfort for a minority of sensitive individuals; such stimuli are called pattern glare stimuli (Nulty, Wilkins, & Williams, 1987; Wilkins, et al., 1984). The archetypal pattern-glare stimuli consist of striped patterns with a spatial frequency of close to three cycles per degree (Wilkins & Evans, 2001). Stimuli of this kind have been implicated in visually induced migraine and, even, epilepsy (Wilkins, et al., 1984; Adjamian, et al., 2004). Accordingly, there is considerable interest in understanding the brain responses to pattern-glare stimuli, since this could indicate the neurological processes that underlie the brain’s susceptibility to headache, migraine, and epilepsy. In this respect, it is believed that pattern-glare stimuli could induce a hyper-excitation of visual processing areas in the brain, a condition that has been implicated in migraine (Welch, D’Andrea, Tepley, Barkley, & Ramadan, 1990).

In a classic MEG study, (Adjamian, et al., 2004) showed that striped patterns at the pattern-glare spatial frequency (∼3 cycles/deg), induced a heightened brain response in the form of increased power at gamma (temporal) frequencies. This gamma response is likely an electrophysiological correlate of hyper-excitation in the frequency domain. More recently, in the time-domain, (Fong, Law, Braithwaite, & Mazaheri, 2020) found differences between migraine sufferers and controls at around 200- and 400-ms post-stimulus onset. Their migraine group showed greater negativity at 200ms for high-frequency gratings (13 c/deg). Indeed, their main findings were on high-frequency gratings, while in contrast, the findings we report occur with clinically relevant, medium-frequency gratings (3 c/deg).

In addition, (Tempesta, Miller, Litvak, Bowman, & Schofield, 2021) presented time-domain findings suggesting that the absence of an N1 (occurring at a time-period between 150 and 200ms post stimulus onset in averaged ERPs) marks the brain’s response to pattern-glare stimuli. Additionally, susceptibility to headache predicted the extent of N1 absence such that participants with increased headache propensity exhibited smaller N1s and thus a more positive-going response. This headache-by-brain-response relationship was observed for repetitions of the pattern-glare stimuli, but not for its first presentation in a sequence of between 6 and 9 repeats (interestingly, (Fong, Law, Braithwaite, & Mazaheri, 2020)’s findings were on the first presentation, perhaps explaining why their findings are different to those we report here, which are on repetitions). This suggests that there may be a habituation component, or failure thereof, to this headache-N1 effect. However, (Tempesta, Miller, Litvak, Bowman, & Schofield, 2021) did not directly determine whether their effect changed across repetitions, e.g. from repetition 2 to repetition 3, 3 to 4 and so on, which would be the definitive test of change through time, whether an increase (sensitisation), or a decrease (habituation). An objective of this paper is to make such a more detailed assessment of the temporal effects (sensitisation / habituation) associated with pattern-glare stimuli across multiple repetitions – which we achieve via a reanalysis of (Tempesta, Miller, Litvak, Bowman, & Schofield, 2021) data.

Habituation (or failure thereof) is a key phenomenon in the study of migraine (and epilepsy), with many studies showing dysfunctional habituation for these conditions (Brazzo, et al., 2011; Coppola, Pierelli, & Schoenen, 2009). Thus, understanding how the brain habituates or sensitises to repeating pattern-glare stimuli is of great interest. For example, if one fully understood how hyper-excitability habituates in the brain, one might be able to use that information to develop therapies, which seek to return the brain to an un-hyper-excited (i.e., habituated) state.

We consider two different patterns of change through time: habituation and sensitization. Both reflect an exponential change through time, with the first matching a brain response that reduces through time; the second one that increases. (The choice of exponential change through time is justified in the Methods.) A key question is the temporal granularity at which any sensitisation or habituation effect operates. For example, it could occur at relatively fine temporal granularity, such as tens of seconds. Alternatively, it may only be observable across minutes or tens of minutes. Fortunately, (Tempesta, Miller, Litvak, Bowman, & Schofield, 2021) presented multiple repeats of each stimulus within each trial with onsets about 4 seconds apart and also divided their experiment into three partitions, each about 12 minutes long thus allowing us to answer our key questions about the granularity of habituation / sensitisation.

We also consider the effects of the state and trait measures introduced by (Tempesta, Miller, Litvak, Bowman, & Schofield, 2021). They conducted a factor analysis on state (discomfort ratings during the experiment) and trait (questionnaire) variables, producing three factors, which were (in order of factor loading): visual stress (trait; sensitivity to visual patterns - particularly stripes), headache (trait; susceptibility to headaches) and discomfort (state; feelings of heightened discomfort when viewing 3 c/deg stimuli during an experiment involving repeated presentation of striped stimuli). These factor scores reflect the susceptibility of each participant to relevant health conditions and symptoms.

**Figure 1:**
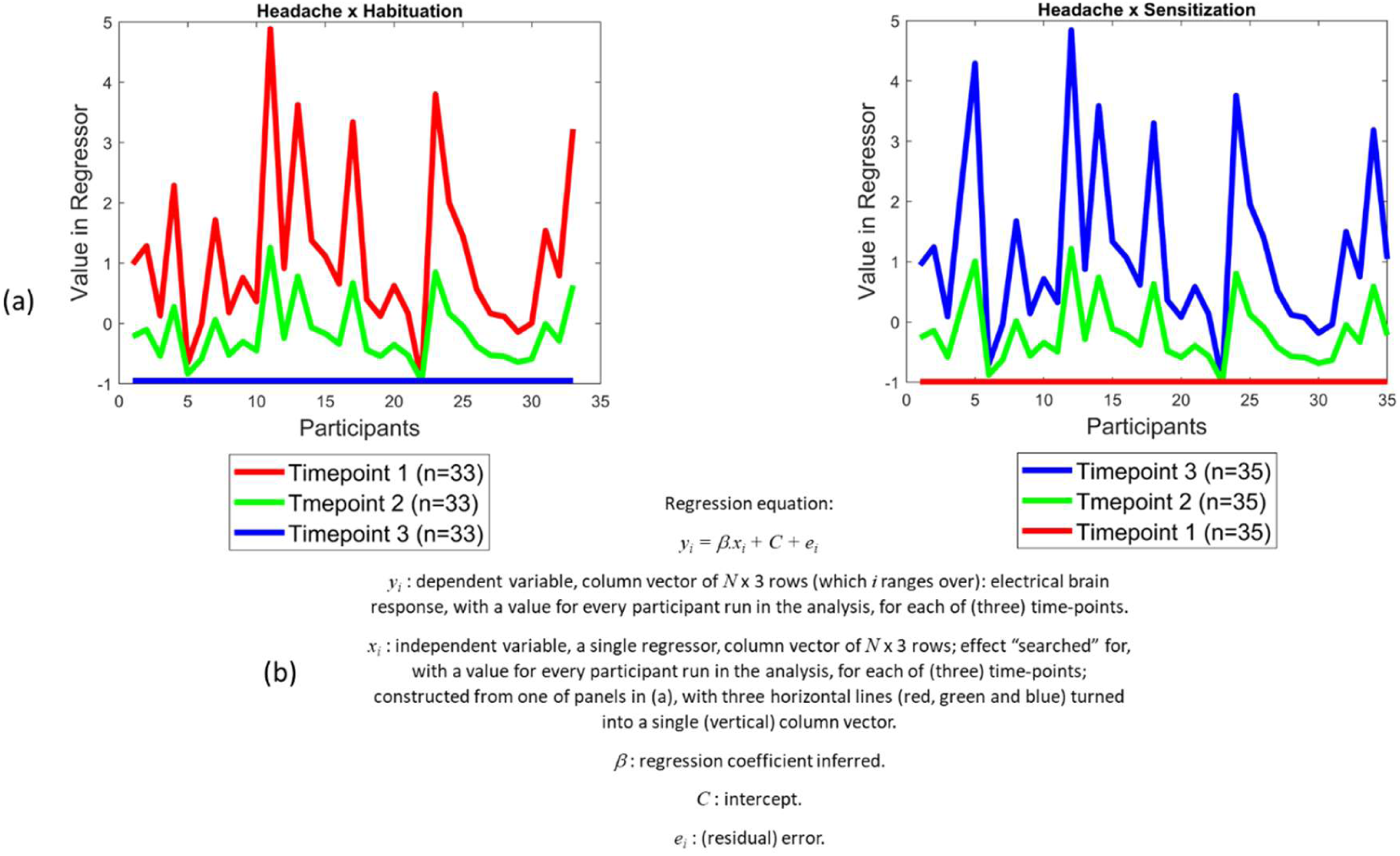
Example of two interaction regressors: (a) shows the Headache x Habituation (left) and Headache x Sensitization (right) interaction regressors. For the habituation regressor, a declining trend is observed through the time-points, representing a habituation effect through time. (For the sensitization regressor, an increasing trend is observed through the time-points.) (b) is the formula (between two panels of (a)) and explanation for the simple linear regression used in the FieldTrip toolbox, since FieldTrip can only perform family-wise error correction with a single continuous regressor (along with the intercept), we have had to build single regressors for all effects.

Based on the above considerations, we formulate our key statistical inference as an interaction between change through time and scores on the three factors. Examples of these interaction effects are shown in Figure Two: InteractionRegressors, with the response to the headache factor changing across three time-points. The first interaction plot (left panel of (a)) models headache-by-habituation. This reflects the intuition that (i) the brain’s response will vary considerably with headache-proneness at the first time point (red line), with those high on the factor showing larger responses, as implied by hyper-excitation, (ii) variability then reduces through time (green line), until (iii) all hyper-excitation has dissipated and all participants exhibit the same response at the final time-point (blue line). Headache-by-Sensitization (right panel of (a)) represents the alternative hypothesis that hyper-excitation emerges during the experiment, leading to high variability in brain response at the final time-point, with those high on the headache factor exhibiting large (hyper-excited) responses. To answer our main question, we test these factor-by-time interactions at the two time-granularities offered by the Tempesta et. al., (2021) data: *fine*, i.e., repetitions within a single trial (with about 4 seconds between onsets) and *coarse*, i.e., partitions of the entire experiment (with each partition lasting 10-12 minutes). We call the fine granularity *onsets* and the coarse granularity *partitions*.

This paper focuses on effects associated with the Headache factor: these being our strongest effects. A companion paper (Dogan, et al., 2023) focuses on the remaining effects, associated with the visual stress and discomfort factors. However, the methods consider, to some extent, all three factors, because we run analyses in which we orthogonalize the headache factor with regard to the other factors. This reflects the fact that they were obtained from the same factor analysis, and we want to guard against benefiting multiple times from the same variability in the data.

To pre-empt our findings, we identify evidence for fine time-granularity (across onsets) sensitization and coarse time-granularity (across partitions) habituation in response to pattern-glare stimuli. Additionally, these effects are modulated by headache, being most clearly evident in those high on the headache factor. This suggests that, in our (non-clinical) population, short time-frame repetition induces hyper-excitation for those susceptible to having headaches, but their brains are able to respond to this hyper-excitation and habituate across the course of the experiment.

## 2. Methods

Our methods are, by necessity, identical to those reported by (Tempesta, Miller, Litvak, Bowman, & Schofield, 2021) whose data we reanalyse. However, for completeness, we present a comprehensive summary here.

### 2.1. Dataset / Participants

Forty participants were initially recruited at the University of Birmingham, all gave informed consent and were compensated with £24 for participating. They had no neurological, psychiatric, or psychological conditions, as well as no history of unconsciousness, convulsions, or epilepsy. Two of the participants were excluded prior to pre-processing, as one left the experiment before completion and the other because of an equipment malfunction. Additionally, during the pre-processing phase we ensure that at least 20% of usable trials exist per condition, per experiment type (Mean/Intercept; Sensitization; Habituation), based on the guidelines by (Luck, 2014). For the Mean/Intercept, Habituation and Sensitization experiments 35, 35 and 36 participants were included respectively. For example, participant A may have at least 20% of usable trials in each group of onsets (2,3; 4,5; 6,7) and therefore would be included in the Sensitization analysis. However, this participant may not have 20% of usable trials in the first partition of the experiment and as a result would be excluded in any Habituation analysis (see Table 1). The study was ethically approved by the Science Technology Engineering and Maths Ethics Committee at the University of Birmingham. Additionally, this study is also in accordance with the relevant regulations set by Scientific Reports.

**Table 1.**
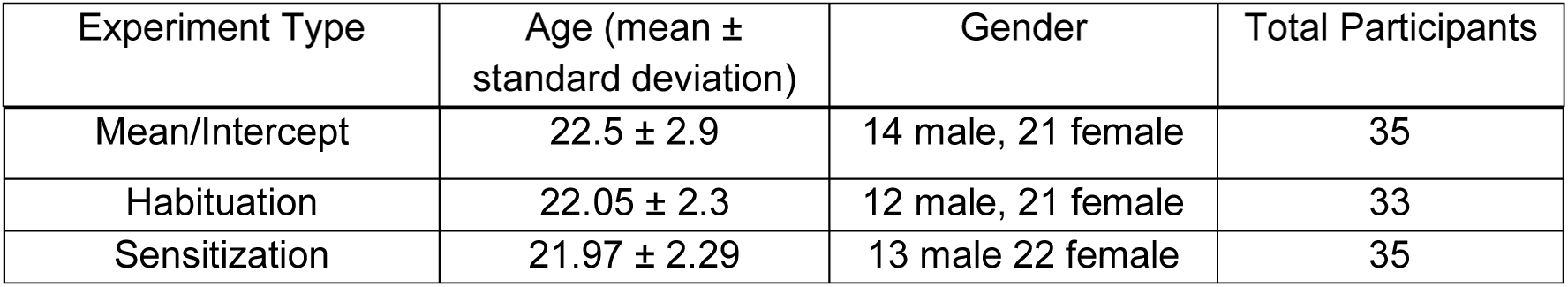
A breakdown of age and gender by experiment type.

### 2.2. Stimuli/Equipment

The stimuli were created with the Psychophysics Toolbox in MATLAB ( (Brainard, 1997; Kleiner, et al., 2007; Pelli, 1997) and were based on those from the Pattern-Glare Test (Wilkins & Evans, 2001)The stimuli were horizontal square-wave gratings at 3 different spatial frequencies (0.37, 3, and 12 c/deg: described as *thick*, *medium*, and *thin* respectively; see Figure 2), and they were displayed at 75% contrast in a circular window with a diameter of 15.2 deg, at a viewing distance of 86 cm. The stimuli were displayed on a 60 Hz Samsung 932BF LCD monitor (Samsung Electronics, Suwon, South Korea) with pixel pitch 0.02 deg/pixel. Each cycle of the 12c/deg grating occupied 4 screen pixels; 3 c/deg, 16 pixels; and 0.37 c/deg, 130 pixels respectively, such that our stimuli were represented without spatial aliasing. Stimuli were calibrated against the monitor’s gamma non-linearity such that the luminance of the grey background matched the mean luminance of the gratings. Pattern 1 (thick) is meant to be a control for low spatial frequency and is not supposed to trigger distortions in most participants. However, it is useful in detecting ‘which patients may be highly suggestible and may respond yes to any question about visual perception distortions’ (Evans BJ, 2008). Pattern 2 (medium) is the only relevant clinical stimulus falling between spatial frequencies 1-4, which are known to elicit migraines and epileptic seizures (Braitwaite, Broglia, Bagshaw, & Wilkins, 2013; Wilkins A. J., 2015). Pattern 3 (thin) is a control for poor convergence and accommodation. Those with poor convergence and/or accommodation will see distortions in this stimulus reflecting optical rather than neurological factors (Conlon E., Lovegrove, Barker, & Chekaluk, 2001).

**Figure 2:**
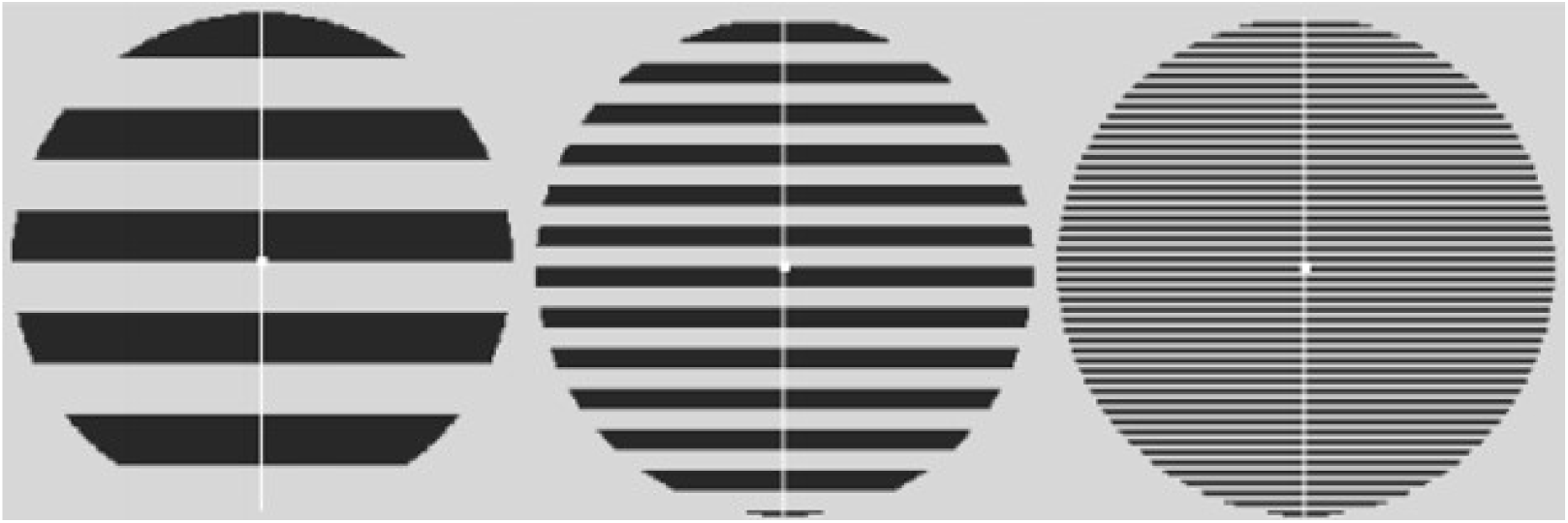
The three types of stimuli (thick, medium, thin) used in the Pattern Glare Test. Note that these images are representative of the stimuli used in the experiment but are rendered here to aid visibility.

The EEG recordings employed a 128-channel BioSemi EEG system and were made in a dark, quiet room.

### 2.3. Questionnaires

For the assessment of headache symptoms, we selected relevant questions from a more general Headache and General Health questionnaire. Thus, we did not use the headache criteria specified by the International Headache Society (Arnold, 2018) to diagnose migraine.

These are criteria for clinical diagnosis and do not provide scale measures of headache proneness. However, the criteria rely heavily on headache intensity, duration, and frequency, and the presence of aura (covering a wide variety of sensory / motor disturbances), all of which were assessed by our questions. For further details of these questions, see Appendix 2 (titled, “Headache Questions”). Questionnaires were also used to assess participants’ tendency to suffer visual stress (Braithwaite, Marchant, Takahashi, Dewe, & Watson, 2015; Conlon E. G., Lovegrove, Chekalu, & Pattison, 1999)

### 2.4. Procedure

Following electrode application, the experiment began with a 5-minute resting period. The main experiment consisted of 3 blocks (partitions) each with 6 trials per stimulus type (thin, medium, thick), totalling 18 trials per stimulus. The trial sequence is shown in Figure 3. Each trial began with a fixation cross displayed for 4 seconds, followed by 7 to 9 onsets of one kind of stimulus for 3 seconds each, and ending with a fixation interval lasting between 1 and 1.4 seconds (varied at random). The stimulus was shown on the screen for the complete 3 second interval, without flicker. After each trial, the participants were asked to rate how comfortable they felt on a scale of 1 (= no discomfort) to 5 (= extreme discomfort), and the number of onsets they observed so that their attentiveness could be estimated. Following each block, the 3 stimuli were presented to the participants consecutively, and they were asked to report the extent to which they had experienced any of the possible pattern-glare symptoms for each one (Wilkins & Evans, 2001). After each block and following the conclusion of the experiment, the participants had a five-minute break during which they were asked to relax and close their eyes. The stimulus order and onsets per trial were counterbalanced between participants.

**Figure 3:**
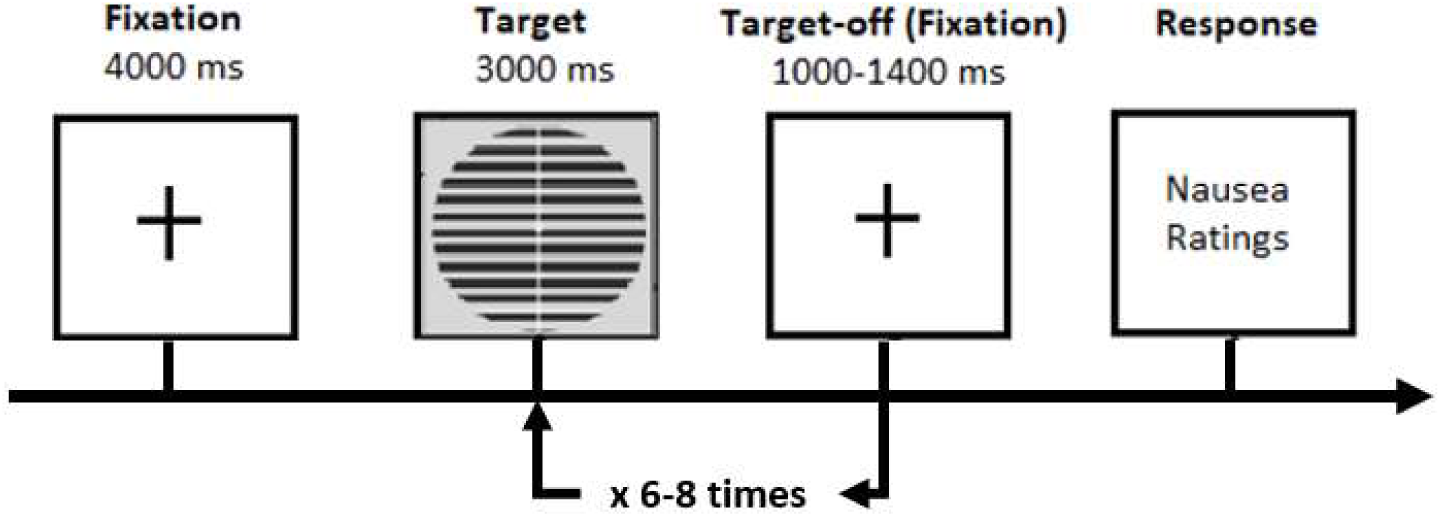
Schematic representation of one trial. This whole sequence was repeated 6 times per stimulus type to complete one block of the experiment.

### 2.5. Factor analysis

Our analysis of the factors followed Tempesta et. al’s analysis and included all 39 participants who completed the study, as factor analysis benefits from larger datasets. Observers vary in their willingness to give high discomfort ratings to stimuli regardless of their form. We allowed for such individual differences by computing a discomfort index (DI) subtracting the mean of the ratings for thick and thin stimuli from that for medium (the latter being the most clinically relevant stimulus). The two visual stress scores (CHi and VDS) were calculated in line with the instructions for each tool. Finally, data for headache frequency, intensity and duration and the experience of sensory aura (see Appendix 2, titled, “Headache Questions”) were extracted.

The seven measures (DI, Chi, VDS, Aura, Headache -frequency, -intensity and -duration) were standardised before factor analysis. A Scree plot identified these factors which, following Varimax rotation, were identified (in eigenvalue order) as visual stress (predominantly a combination of CHi, VDS and aura), headache (frequency, intensity, and duration) and discomfort (DI). Initial factor scores were computed using the regression method from coefficients shown in Appendix 3 (titled “Factor Analysis”), where we also describe the factor analysis in more detail. Further analysis investigated the reliability of applying dimensionality reduction techniques (PCA; Factor Analysis; Averaging) to our behavioural data. We concluded that, Factor Analysis is a reliable decomposition method in the context of our data. Please refer to Appendix 4 (titled “Justification of factor analysis decomposition”).

The factor structure described above is unsurprising given the variables included, but the analysis was also intended to provide orthogonal factors. When we further reduced the number of participants, as a result of artefact rejection, our factors were no longer fully orthogonal, although this loss of orthogonality was small. Nonetheless, departing from (Tempesta, Miller, Litvak, Bowman, & Schofield, 2021), we orthogonalized headache and discomfort with respect to visual stress (the strongest factor) on this smaller number of participants. This was done using the Gram–Schmidt algorithm (Arfken, 1985).

### 2.6. EEG pre-processing

Our EEG processing diverges considerably from that of (Tempesta, Miller, Litvak, Bowman, & Schofield, 2021) although in common with them we down sampled the EEG data from a sampling rate of 2048 to 512 Hz using the Biosemi toolbox. Eye-blink artefacts were removed using independent component analysis (ICA), with ICA components associated with eye blinks removed and the dataset reconstructed.

The FieldTrip toolbox (Oostenveld, Fries, Maris, & Schoffelen, 2011); version 20210807, was then used for the following pre-processing and analysis. EEGs were band-pass filtered with a FIR filter using a range of 0.1 - 30 Hz using a Hanning window. Data for each onset were epoched between -200 and 1200ms relative to stimulus onset, referenced to the average of all electrodes and baseline corrected based on the 200ms period prior to stimulus onset. Thresholding was then applied with a 100μV threshold in both the positive and negative directions, thus removing any large artefacts present in the data. Participants were rejected if after thresholding less than 20% of the original trial count remained for any one of the three conditions (Thick, Medium, and Thin), or experiment types (Mean/Intercept, Habituation and Sensitization).

The stimuli were equally probable at the first onset; as a result, onset 1 was not considered in the analysis because it could contain response information reflecting surprise. In contrast, the stimuli presented from onset 2 onwards (within each trial) were completely predictable.

As well as the Factors dimension (which, as previously discussed, has three levels: Visual Stress, Headache and Discomfort), we had a dimension reflecting change through time. For this dimension, we considered two different patterns of change through time: *habituation* (a reducing brain response through time) and *sensitization* (an increasing brain response through time). These two different patterns of change through time, were applied to two different time granularities (which give the dependent variable for our regressions). The first of these reflected fine-grained changes through time, by grouping Onsets 2-7 into three bins (2,3; 4,5 and 6,7). The second reflected coarse-grained change through time, i.e. across the course of the entire experiment, which was investigated by dividing the experiment into its 3 natural partitions (P1, P2, and P3), representing each block of the experiment for onsets 2:8. Finally, the three-way-interaction (onsets 2:3, 4:5, 6:7 across partitions P1, P2 and P3) combines both habituation and sensitization into one interaction regressor, with the key hypothesis representing a decrease in sensitization through the course of the experiment. All remaining contrasts, such as the Mean/Intercept and Analysis on Factors, were performed on onsets 2:8.

### 2.7. Design Matrices and Mass-univariate Analysis

A Mass Univariate Analysis (MUA) was conducted in FieldTrip (Oostenveld, Fries, Maris, & Schoffelen, 2011) on our ERP data, using FieldTrip’s cluster-based permutation test method. The significance probabilities of the permutation tests were calculated using the Monte Carlo method, and all tests were run for 25,000 permutations.

The MUA was performed on what we name the Pattern Glare Index (PGI). This index enabled us to focus our study on where the clinically relevant medium stimulus exhibited more extreme responses than the thin and thick stimuli. The equation used for its calculation was the following:

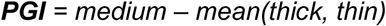

The analysis included regressing the data onto the following predictor variables:

1. The intercept of the regression model was a predictor variable, with data collapsed across onsets 2-8 and across the entire experiment. Since all regressors entered into the regression model are mean-centred, the intercept becomes the mean of the basic stimulus-effect on the PGI, which identifies data points in which medium is extreme relative to thick and thin.

2. The factor scores derived from the factor analysis were used as regressor predictors, with data collapsed across onsets 2-8 and across the entire experiment. The factors were identified as follows: *visual stress, headache,* and *discomfort*.

3. Continuous regressors representing the change through time were used as predictors. We considered two different profiles of change through time: *habituation* and *sensitisation*. Habituation reflects an exponential decrease, whereas sensitization reflects an exponential increase, given in the following form:

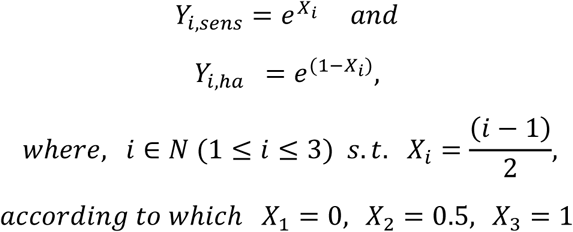

Thus, we raise *e* to the power of a linear decrease in the habituation regressors, and a linear increase in the sensitization regressors. The resulting exponential decrease or increase was selected as neural responses are typically better described by exponential changes, for example, exponential decay is common in biology and neural dynamics (Trappenberg, 2009), reflecting the non-linearity of neuron firing. The regressors, after mean-centring, are visualised in Figure 4, where *time*_*point i* = *Y*_*i,hab*_ on the left-hand side and *time*_*point i* = *Y*_*i,sens*_ on the right-hand side.

**Figure 4:**
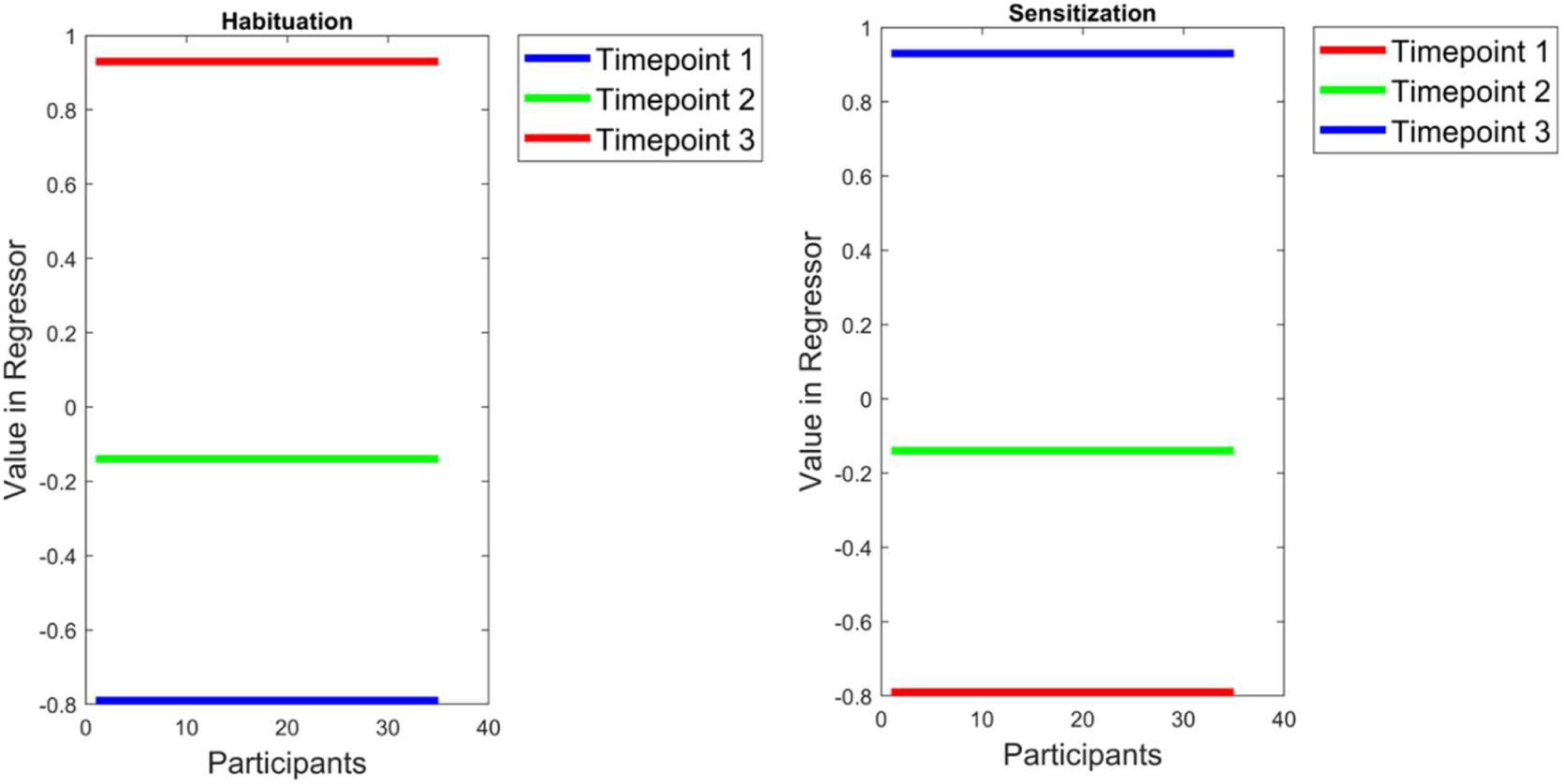
(Left) A regressor representing habituation through time, (Right) a sensitization regressor representing sensitization through time. The x-axis represents the participants and the y-axis the design matrix value.

4. (Unorthogonalized) Factor x Time interaction regressors were entered into regression models, where Factor is Headache and Time could be Habituation or Sensitization.

The habituation and sensitization regressors resulted from the following procedure: firstly, for both types of regressors, we generated a baseline-shifted form of each factor, to have a completely positive regressor. This effectively assumes that participants that scored low on each factor would not habituate or sensitize. We did this by subtracting the minimum (i.e., most extreme negative) score from each participant’s score. Then, the baseline-shifted scores were multiplied with the exponential decrease in the habituation case, and the exponential increase in the sensitization case. Lastly, in both cases, the regressor was mean centred.

The resulting Headache x Habituation, as well as Headache x Sensitization interaction regressors are shown in *Figure 1*. Thus, in the habituation case, the resulting interaction regressor reflects the intuition that the greatest differences in brain responses should be observed in the first time-period, with the PGI response being much larger as the score on the factor increases. Then, habituation should mean that differences in brain responses reduce on the second time-period and disappear by the third, leaving all participants with the same low response reflecting the extinguishing of hyper-excitation. (This is not the pure interaction between a factor and habituation since that would not fully reflect our intuition of habituation. In particular, since there is a reduction in hyper-excitation across the course of the experiment, the average response is also reducing through the partitions. However, for presentational ease we still use the term interaction.) Conversely, in the case of sensitization, the resulting interaction regressor expresses no factor differences at the first time-point, more at the second and finally, the biggest differences at the last.

These two-way interaction regressors, Factor x Time (where Factor is Headache and Time could be Habituation or Sensitization), were applied over both time granularities: Onsets and Partitions.

5. Orthogonalized Factor x Time interaction regressors were entered into regression models, where Factor is Headache and Time could be Habituation or Sensitization.

The orthogonalization was performed on the baseline-shifted and mean-centred versions of the regressors, which for Headache x Habituation is depicted in *Figure 1* (left side). This was done as the interaction regressors were not orthogonal. The other two interaction regressors were orthogonalized with respect to the Visual Stress x Habituation interaction regressor and themselves, using the Gram–Schmidt (Arfken, 1985) process. This sequence of orthogonalizations was chosen because Visual Stress was the factor that obtained the strongest loading in our factor analysis and thus would naturally be preserved unchanged by the orthogonalization.

6. Finally, a three-way interaction regressor was entered into the MUA. This regressor represents a Factor x Habituation {for Partitions} x Sensitization {for Onsets} for Headache.

The three-way interaction regressor consists of both the Factor x Habituation {for Partitions} and Factor x Sensitization {for Onsets} factor scores described under point 5. This regressor consists of the following: The Factor x Habituation {for Partitions} contains three partitions with each partition consisting of all subjects (3 partitions x 32 subjects), within each partition, three groups of onsets exist, 2;3, 4;5 and 6;7. This results in 9 groups for the three-way-interaction. Once the regressor is constructed, the procedure described under point 5 is applied to this regressor.

The three-way interaction regressor builds on both the two-way interaction regressors {factor x habituation for partitions} and {factor x sensitization for onsets}. The intuition reflects the hypothesis that, during the course of the experiment, participants exhibit a decreased sensitization through the onsets. Thus, at the beginning of the experiment (partition one), hyper-excitation increases substantially through the onsets and the extent of that increase is dependent upon participants’ factor score (higher on factor, more sensitisation). This factor by sensitization pattern reduces in partition 2 and then disappears by the end of the experiment (partition 3). Figure 5 depicts the three-way interaction regressor for the Headache factor. Thus, what we see in Partition 1 is similar to what we would observe for the two-way interaction of factor by sensitization for the Onsets. Then, Partition 2 is similar, but with the factor by sensitization for Onsets pattern “turned-down”, i.e., less difference across Onsets (the sensitization) and across participants (the factor). Finally, Partition 3 reflects a complete absence of the factor by sensitization for Onsets pattern, i.e., nothing changes across Onsets or participants (the factor). Additionally, since there is a reduction in hyper-excitation across the course of the experiment, the average response also decreases through the Partitions.

**Figure 5:**
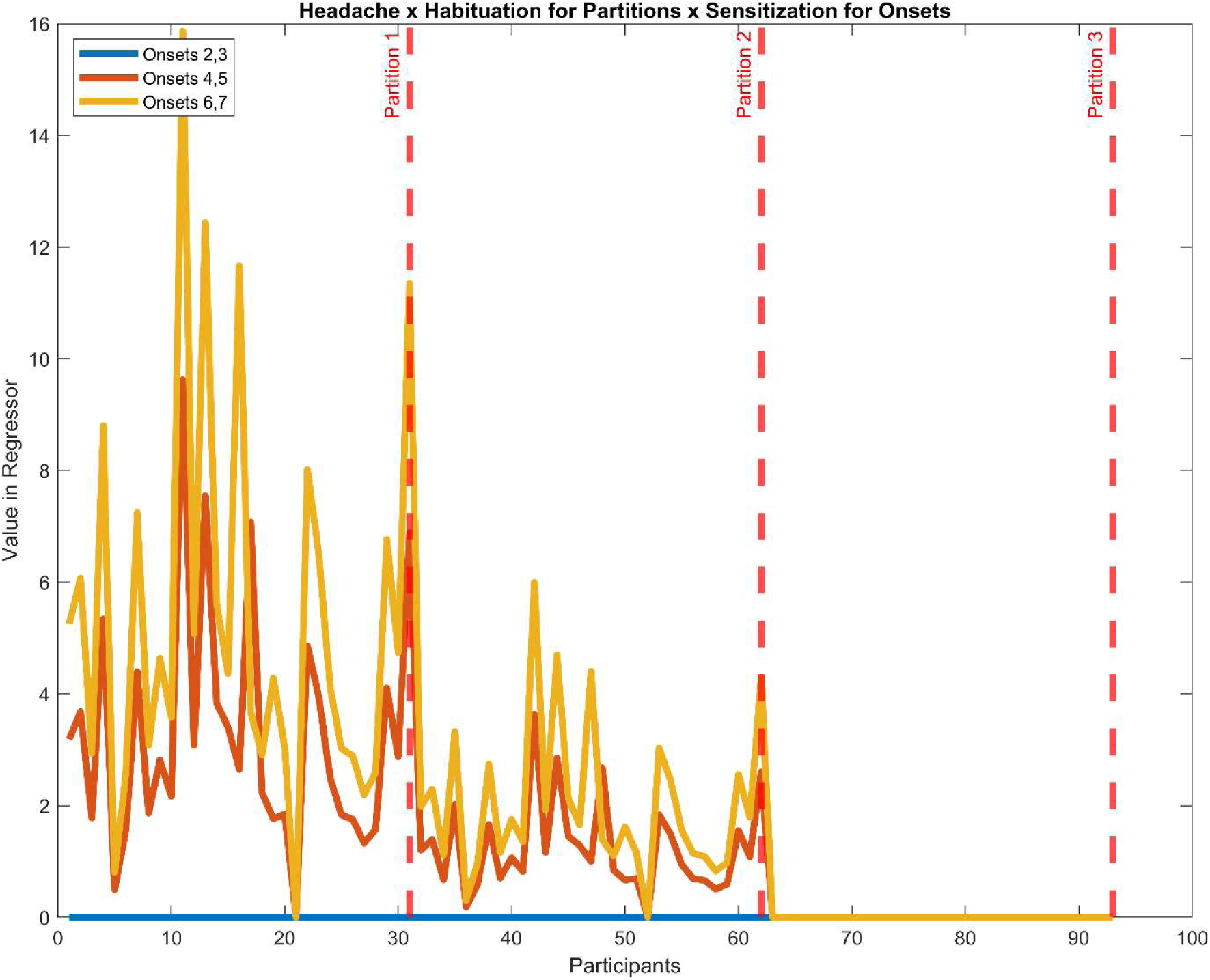
Example three-way-interaction regressor for the Headache x Habituation for Partitions x Sensitization for Onsets effect: The x-axis represents participants, repeated for each partition. The y-axis indicates the design matrix value assigned. Onsets are indicated by the blue, orange, and yellow lines (2,3; 4,5; 6,7) with partitions segmented by the red dashed lines.

FieldTrip, the MUA analysis software used, only enables regressions to be formed with an intercept and one other regressor. This is due to the technical difficulties associated with performing the cluster inference permutation test over complex design matrices. Consequently, one is forced to run multiple separate regressions, with different regressors. This raises the possibility that the same variability in the electrophysiological data could “count multiple times”, i.e., contribute to making multiple regression coefficients extreme (relative to 0). Accordingly, we explicitly highlight any inferences that we make that could be inflated in this way as exploratory findings. This particularly arises when we focus on the un-orthogonalized headache-by-time interaction regressors (see point 4 above), which, as we have indicated, are collinear, with the same regressors for other factors.

The parameters inferred for regressors were statistically examined through two-tailed one-sample t-tests (testing the difference of a single regression coefficient to zero) at the voxel level and we searched for significant clusters (FWE-corrected with p < 0.05, cluster-level), with a cluster-forming threshold of 2.5% (i.e., alpha=0.025).

### 2.8. Regions of Interest

Similarly, to (Tempesta, Miller, Litvak, Bowman, & Schofield, 2021), we considered the evoked transients observed at posterior electrodes, arising from stimulus-onset. Following (Tempesta, Miller, Litvak, Bowman, & Schofield, 2021), we limited our analyses to a space-time region of interest (ROI) centred around the posterior electrodes and seeded by an overall temporal analysis (*bounding*) window, determined by deviations from baseline in the time-series. This was a two-stage process: calculation of initial bounding window followed by identification of final ROI, within this bounding window. This final ROI is three dimensional, i.e. a region in time and (two dimensions of) space.

We calculated the bounding window as follows. We sought to capture the time period where the ERP was significantly different from baseline by focussing on the *aggregated average* (Bowman, Brooks, Hajilou, Zoumpoulaki, & Litvak, 2020; Brooks, Zoumpoulaki, & Bowman, 2017) across all stimulus conditions (three spatial frequencies, Onsets 2-8, and three partitions). This approach does not inflate the false-positive rate as it does not reflect stimulus differences or changes over time (i.e., over partitions or onsets), since it is an orthogonal contrast (strictly, parametrically contrast orthogonal) to the contrasts we are interested in here (Bowman, Brooks, Hajilou, Zoumpoulaki, & Litvak, 2020; Brooks, Zoumpoulaki, & Bowman, 2017).

This aggregated average did not settle back to baseline as would normally be expected, but rather fell to a constant Direct Current (DC) level, which was maintained until stimulus offset (the post-transient baseline). To address this, we captured the period from the initial (statistically significant) deviation from zero to when the aggregated average fell to the point of not being statistically different from the DC level, as calculated over a 400s period well after the end of the stimulus transients. This was done at electrode Oz (A32), since it is the electrode that would, a priori, be most expected to show visually evoked transients. We then calculated confidence intervals across participants, weighted by the number of valid trials and compared to the pre- and post-transient baselines (as appropriate). The equations for the weighted CI at each time point are given by:

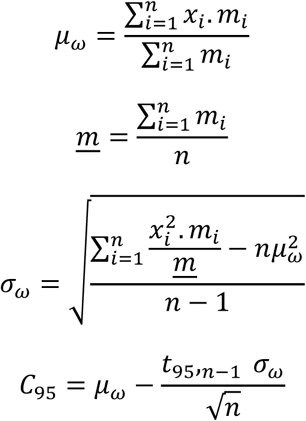

where *μ*_*w*_ is the weighted mean, <u>*m*</u> the mean number of trials per participant, *σ*_*w*_ the weighted standard deviation, *C*_95_ the confidence interval, *n* the number of participants, *t*_95,*n*-1_ = 1.7 the critical t value for a one-tailed 95% confidence interval, *x*_i_ the value of the ERP for the ith participant, and *m*_i_ is the number of valid trials for that participant. The lower CIs were compared to zero (pre-transient baseline) at the start of the grand-average trace and to the DC level (post-transient baseline) at the end of the trace.

The resulting bounding window (56-256ms) was used to seed our final ROI based on the mean/intercept effect (see Results). All (non-intercept) regressors are mean-centred, ensuring that the intercept of the regression becomes the grand mean and is orthogonal to all (non-intercept) regressors. This ensures that (non-intercept) regressors are parametrically contrast orthogonal to the mean/intercept regressor in the sense highlighted in (Bowman et al, 2020), and ROI selection on the mean/intercept does not inflate the type-1 error rate when testing the (non-intercept) regressors in that ROI.

We used one-sample t-tests to demonstrate that individual regression coefficients (for our factors, time, factors-by-time, factors-by-time-for-onsets-and-partitions, and intercept regressors) are statistically different from zero. All analyses were run two-tailed.

Additionally, to enable reproducibility of our results, we have included an additional file containing the ROI used in this analysis.

### 2.9. Data Visualization

MUA finds statistically significant relationships between regressors and space-time ERP maps. This approach does not provide a complete visualisation: for example, it does not show how particular differences between stimuli underlie an effect. We included ERP time-series plots to provide this extra information, placing our participants into two groups for each factor based on median splits of the factor scores.

A further analysis was developed to select the appropriate channel for visualisation purposes. As part of the MUA, FieldTrip reports a collection of statistics over a 3D matrix, two representing space and one time, including significance levels (p), and t-statistics. The electrode used for visualising ERPs was selected by computing the cumulative count of significant samples / voxels for each channel, i.e. where the significance level was ≤2.5% (i.e., probability of 0.025) at that point in time and space. The electrode with the highest count in a significant cluster was then selected for visualisation.

Following the above electrode selection, 95% confidence intervals (CI) for each time series were computed using bootstrapping. Firstly, *N* participant ERPs are sampled with replacement, where *N* is equal to the number of participants. Sampling with replacement allows participants to be selected multiple times, or not at all, in each sample thus introducing variability between the samples based on the variation in the original dataset. The grand average of these bootstrapped ERPs for each condition were then computed and stored. This process is repeated 3,000 times to create a distribution over participants bootstrapped grand-averaged ERPs. Finally, for each timepoint in the time series, 2.5% and 97.5% percentiles were calculated based on the bootstrapped distribution to create confidence intervals around the ERPs.

## 3. Results

The results that follow are separated into five main components: regressing the PGI onto 1) the Mean/Intercept; 2) the Factors; 3) the Factors x Time for Partitions interaction regressors; 4) the Factors x Time for Onsets interaction regressors; and 5) Factors x Time for Partitions x Time for Onsets interaction regressors. The last of these was conducted because the Headache x Habituation for Partitions and the Headache x Sensitization for Onsets were themselves significant suggesting a specific three-way interaction. Thus, we only ran the corresponding three-way interaction: Headache x Habituation for Partitions x Sensitization for Onsets; no other three-way interaction was tested. (A 6th pure change through time regressor was also analysed but is reported in a companion paper.)

We are most interested in the positive going effects at the occipital electrodes, so the following figures are targeted at those effects. We adopt this focus because the mean/intercept analysis revealed a positive going effect in the occipital area, i.e., over visual cortex, suggesting that the hyper-excitability generated by the medium stimulus manifests as a more extreme response in a positive going direction over visual cortex. Since all our further analyses are carried out in the ROI identified from the mean/intercept, the association of medium with a more extreme positive going pattern is carried over to these further contrasts, i.e., on Factors; Factors x Time for Partitions; Factors x Time for Onsets; and Factors x Time for Partitions x Time for Onsets.

### 3.1. Mean/Intercept Effect

MUA found statistically significant clusters for our mean/intercept effect across the whole scalp and through time (within our bounding window from 56-256ms: see methods). This was our statistically strongest effect, and it was mostly present in the occipital area of the scalp, as expected.

In Figure 6, we can see three different panels capturing the effects between 56-256ms. Figure 6(a) shows the largest, most positive-going cluster, for the PGI, as a percentage of the entire scalp. The effect is largest at approximately 160-190ms, accounting for just over 20% of the volume.

**Figure 6:**
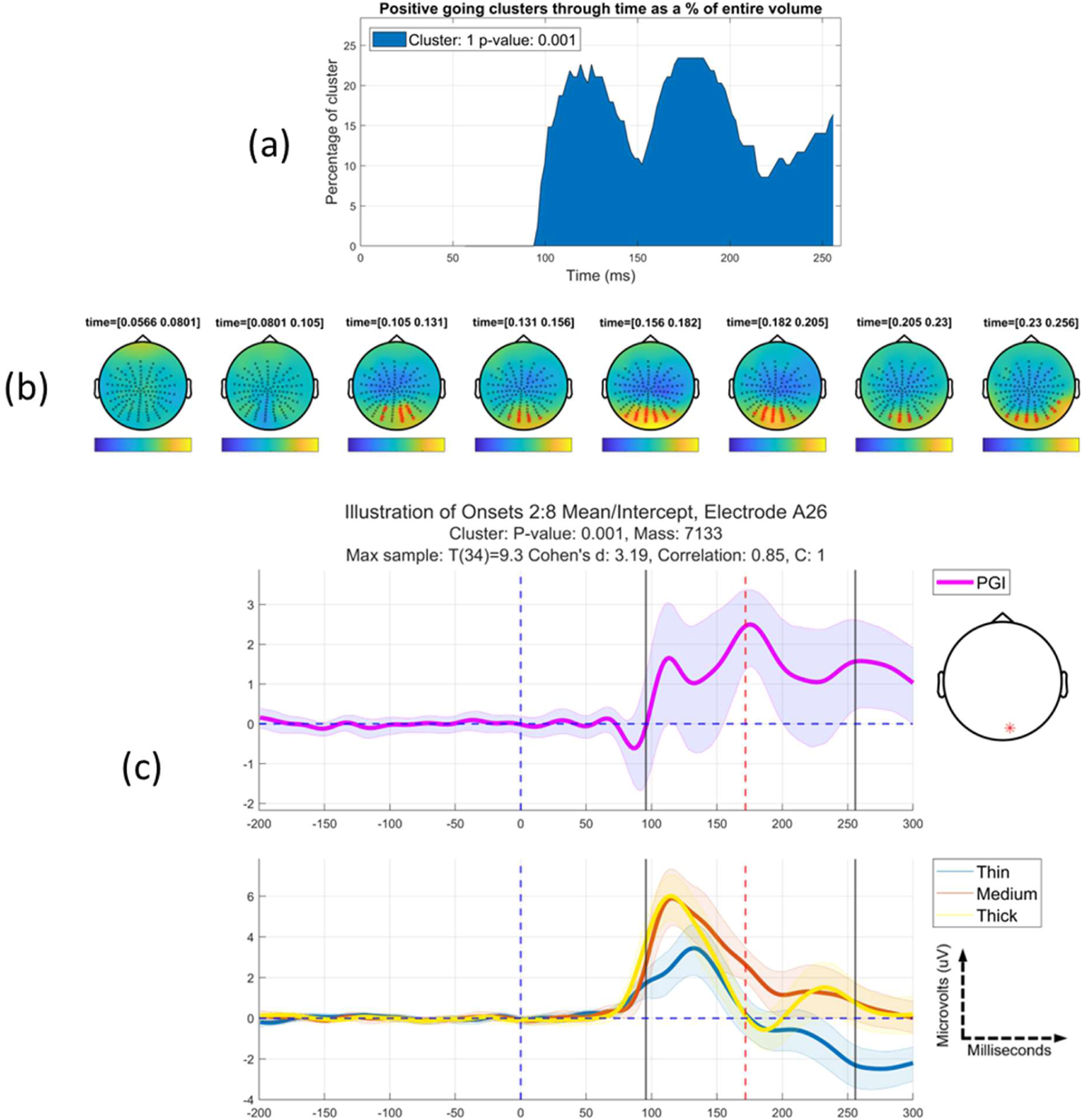
Mean/Intercept effect: (a) The most positive going cluster, for the PGI, through time as a proportion of the entire scalp. (b) Topographic maps through bounding time window, 56-256ms after stimulus onset, with intervals of 25ms. A very strong positive-going cluster is observed at posterior electrodes, while an associated central negative-going cluster is also present, likely reflecting the opposite side of the driving dipole. (c) Grand average time-series of the PGI and individual conditions at the electrode that is the most continuously significant through time. The shaded regions represent 95% confidence intervals for each time-series. The black lines indicate the start and end of this positive (most significant) cluster through time, with the red dashed line indicating the peak of the effect. The clinically relevant stimulus (medium) is clearly stronger than the mean of the control stimuli (thick and thin) for an extended time period, reflecting, we would argue, a time-domain correlate of pattern-glare induced hyper-excitation.

Figure 6(b) shows the scalp topography maps through time during our bounding window 56-256ms after stimulus presentation. The topographic plots represent the unthresholded t-values at each time-space sample, with a scale adjusted based on the maximum t-value present in the resulting statistical test. Electrodes belonging to the significant (family-wise error, FWE corrected) cluster are highlighted in red. These indicate that the effect is mostly at the posterior of the scalp, with the effect lasting between approximately 100 and 256ms. The cluster has a p-value ≤ 0.001 (FWE-corrected, cluster-forming threshold 0.025), with the peak of the effect at 173ms. Figure 6(c) shows the grand average ERPs for the most continuously significant electrode in the cluster. These time-series are split into two subplots: the top panel displays the PGI, with the panel below showing the individual stimulus conditions (thin, medium, and thick stripes) that contribute to the PGI. The shaded areas represent the 95% confidence intervals. The black vertical lines represent the start and end of the most positive going cluster through time, with the red dashed vertical line representing the peak of the effect.

Table 2 summarises the significant clusters revealed by the MUA for the Mean/Intercept regressor. Both negative and positive tails are included, and the information displayed are the results of the significance test (i.e., probability of arising under the null), the cluster’s most sustained electrode, the time of this electrode’s peak and t-value of this electrode’s peak; and the sum of the t-values across the whole cluster.

**Table 2.** Collective results for both positive (+1) and negative (-1) tails of the mean/intercept effect. For both the positive and negative going clusters, the following values are presented: the significance probability (p-value, FWE-corrected); the electrode where the effect is most sustained through time; the time of the peak for the most sustained electrode; the t-value at that peak; and the summed t-values of the significant cluster. Clusters are ordered from most to least significant; * indicates p≤0.05.

| *Mean/Intercept* |  |  |
| --- | --- | --- |
| *Statistic* | *Positive Cluster 1* | *Negative Cluster 1* |
| *p* | $\leq 0.001^*$ | $\leq 0.001^*$ |
| *Most sustained electrode* | A26 | C24 |
| *Time of peak of most sustained* | 173ms | 174ms |
| *Max T(34)-value* | 9.3 | 6.8 |
| *Cluster Stat* | 7,133 | 10,155 |

Consistent with Figure 6, Table 2 shows two large clusters, the positive-going posterior one we focus on in Figure 6 and the second negative-going more central cluster, seen in Figure 6(b) (p≤0.001; FWE-corrected; cluster-forming threshold 0.025). These suggest a sustained hyper-excitation response to the (aggravating) medium spatial frequency stimulus.

The posterior positive going cluster that we observe here becomes the ROI that we use for all future analyses. We chose to focus on this cluster, since it most directly conformed with our prior hypotheses. Furthermore, although also large, the more central negative-going cluster looks to be the opposite side of the electrical dipole generating the positive cluster. (Indeed, such polarity reversals on the scalp are typically observed when an average reference is taken.) Consistent with this, the size of the negative-going cluster waxes and wanes in synchrony with the positive-going cluster, suggesting that the two effects are strongly negatively correlated. This, in turn, suggests that no more substantive information would be gleaned from also exploring the negative-going effect further.

### 3.2. Analysis on Headache Factor

In this section, we present analysis for the Headache factor for all participants with cluster inference being performed within the ROI identified by the mean/intercept contrast.

One significant positive-going cluster was identified for the Headache factor (see Figure 7) over occipital lobe within the mean/intercept ROI. The effect spanned 125-192ms. The (FWE-corrected) *p-*value is 0.046, with the peak of the effect found at 152ms with a *t(33)*-value of 3.83 (Table 3).

**Figure 7:**
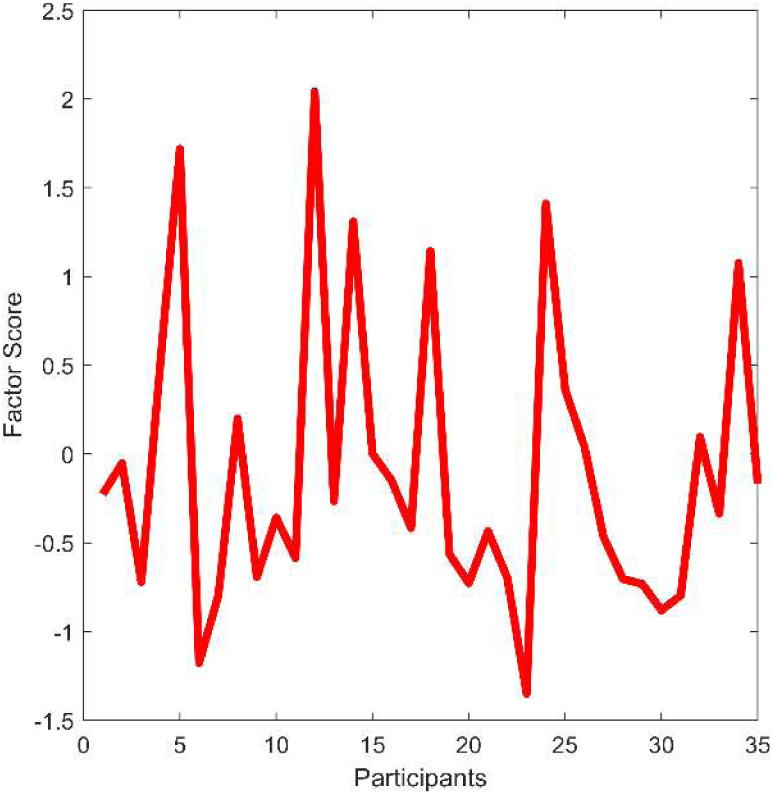
Headache factor regressor. The x-axis represents the participants and the y-axis the design matrix value assigned to each one.

**Table 3.**
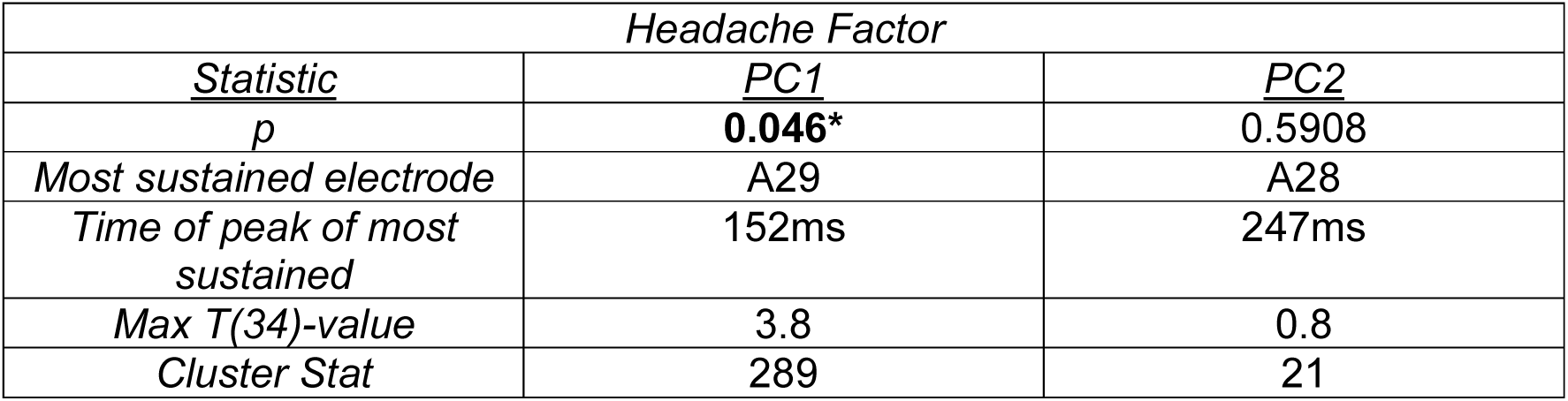
Collective results for the positive (+1) tail for the analysis on the headache factor. For each effect that crossed the cluster forming threshold, the following values are presented: the significance probability (p-value, FWE-corrected); the electrode where the effect is most sustained through time; the time of the peak for the most sustained electrode; the t-value at that peak; and the summed t-values of the significant cluster (i.e., the Cluster stat). Clusters are ordered (left to right) from most to least significant; PC = positive cluster found within the ROI. * Indicates p≤0.05.

Figure 8(b) shows the scalp topography maps through time during our bounding window (56-256ms) for the significant cluster. As seen in both panels, HeadacheFactor(a) and HeadacheFactor(b), the effect starts at 125ms lasting until 192ms, centred over the occipital lobe with the peak electrode being A29. Figure 8(c) displays the grand average ERPs for the most continuously significant electrode in the cluster divided into high (right column) and low (left column) according to a median split on the Headache factor scores.

**Figure 8:**
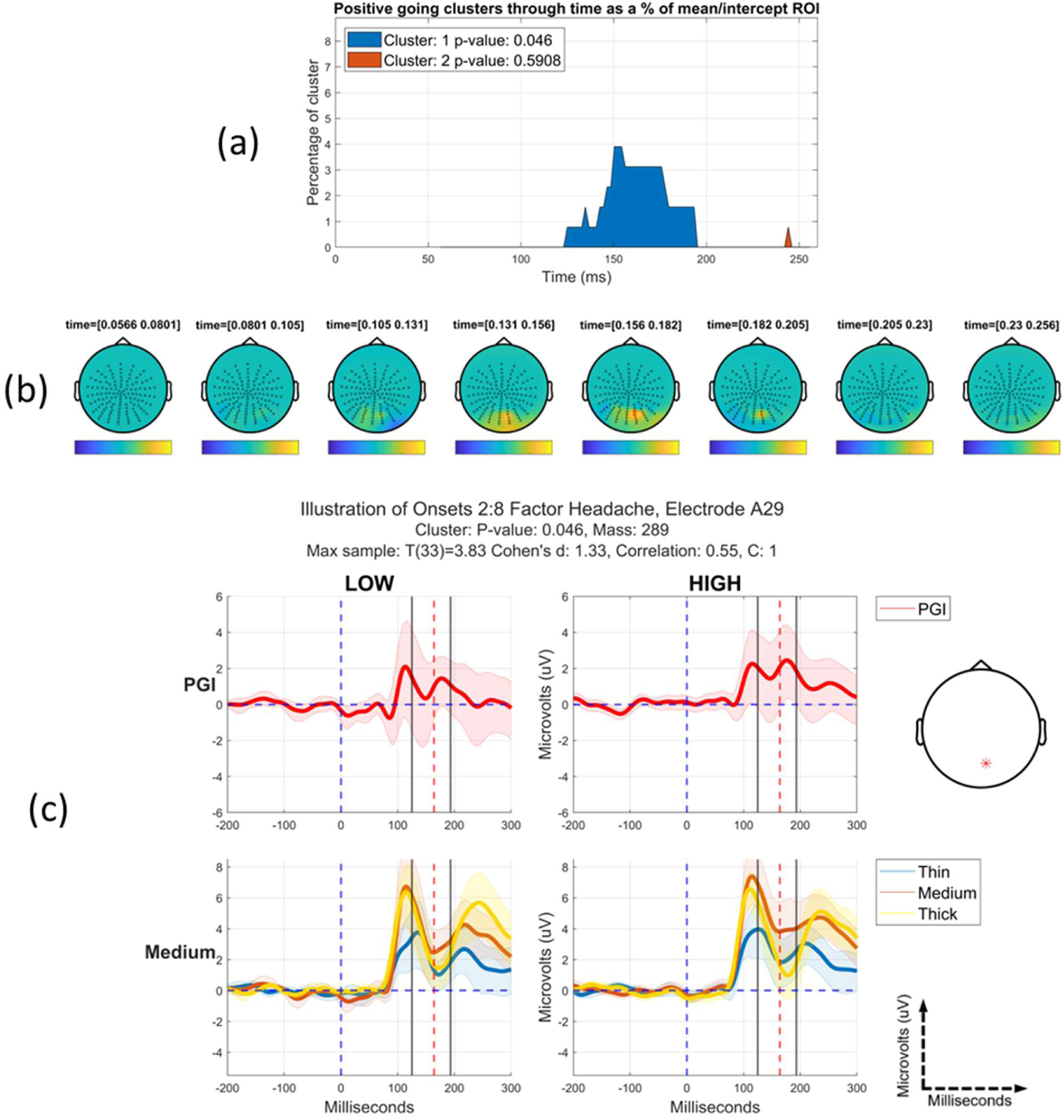
(a) The most positive going clusters through time as a proportion of the mean/intercept region-of-interest. (b) Topographic maps through bounding time window, 56-256ms after stimulus onset, with time step between each map approximately 25ms. A positive-going cluster is observed at posterior electrodes. (c) Grand average time-series of the PGI (top-row) and individual conditions (bottom-row) at the electrode that is the most continuously significant through time. Left and right columns reflect a median split on the Headache factor scores into Low and High groups. The shaded regions represent 95% confidence intervals for each time-series. The black lines indicate the start and end of the most significant cluster through time, with the red dashed line indicating the peak of the effect. During the window of significance, the clinically relevant stimulus (medium) is substantially stronger than the mean of the control stimuli (thick and thin) for the high group and much less so for the low group, with this effect lasting for an extended time period. This suggests a time-domain correlate of pattern-glare induced hyper-excitation that is sensitive to susceptibility to headache.

This effect suggests a time-domain correlate of pattern-glare induced hyper-excitation that is sensitive to susceptibility to headache.

### 3.3. Two-way Interactions: Factor-by-Time

These two-way interactions are key contrasts, since they encapsulate how our clinical-condition variable (i.e., headache) modulates change through time (habituation or sensitization), and the granularity of time (Onsets or Partitions) at which that modulation manifests. Across the headache factor considered here and other factors considered in the companion paper, the only two-way interaction effects that survived FWE correction were Factor-by-Habituation for Partitions and Factor-by-Sensitization for Onsets. For the headache factor, we focus here first on habituation (see section 3.3.1) and then on sensitisation (3.3.2), with all other effects summarised in section 3.5, Table 8.

For each time pattern, there is collinearity between the three relevant interaction regressors. For example, considering habituation across the partitions, after combining the factor scores with the Habituation effect there is strong collinearity, which is largely driven by the Habituation effect being present in all the interaction regressors. Consequently, we present our two-way interaction results in two parts: 1) basic effects, which are on orthogonalized regressors; 2) effects on unorthogonalized regressors, which serve as exploratory findings. We only present statistically significant results here. Close to-, but not statistically significant, effects are presented in Appendix 5. Additionally, effects that come out in both the orthogonalized and non-orthogonalized analyses can be considered strong. This is because the former of these is the statistically most principled, but the orthogonalization process can distort the regressor. Thus, an effect that also comes out in the absence of orthogonalization is assured to also conform to the intuition we have for our two-way regressors.

#### 3.3.1. Non-orthogonalized Headache x Habituation for Partitions (Exploratory)

In this section, we present the non-orthogonalized, exploratory Headache x Habituation for Partitions interaction regressor. The interaction for the Orthogonalized factor was not significant (cluster p=0.0663) but showed some similar effects to those presented below (see Appendix 5, section “Orthogonalized Headache x Habituation for Partitions” for orthogonalized version). MUA identified one statistically significant positive-going cluster when regressing the Headache x Habituation regressor onto the Partitions (depicted in Figure 9). The (FWE-corrected) *p-*value is 0.0033 with the peak of the effect found at 133ms with t(97) of 3.91 (Table 4).

**Figure 9:**
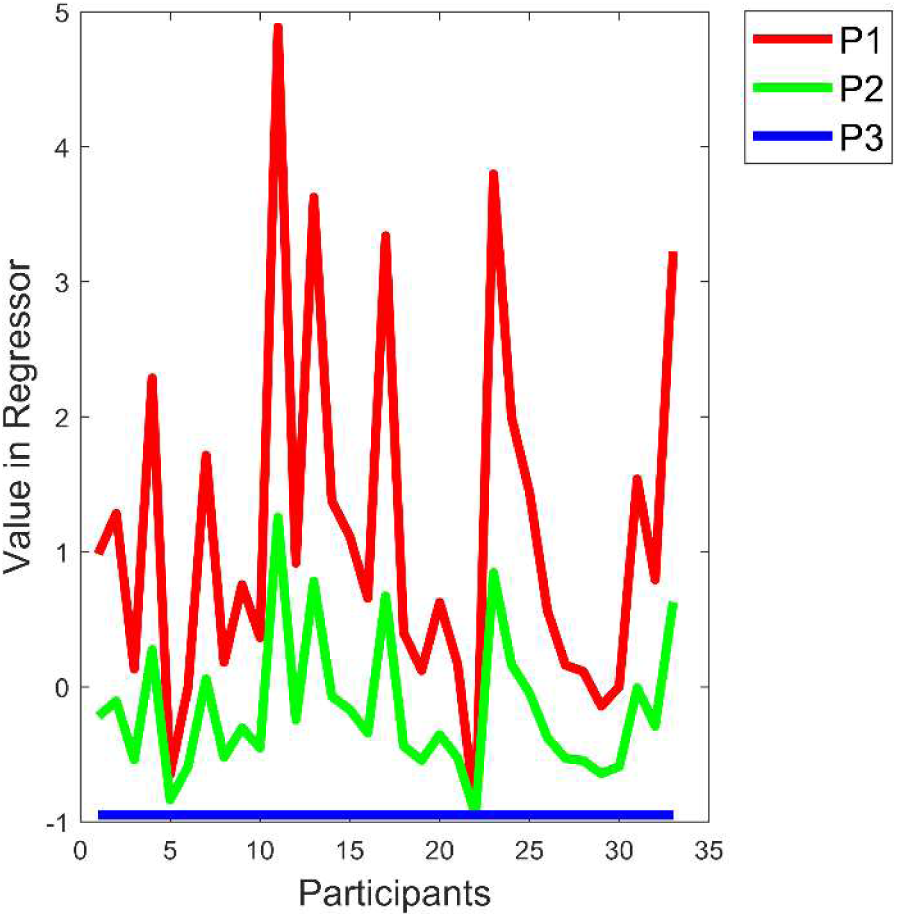
Headache x Habituation for Partitions interaction regressor. The x-axis represents the participants and the y-axis the design matrix value assigned to them for each partition. Partitions are indicated by P1, P2 and P3.

**Table 4.** Collective results for positive (+1) tail of the Headache x Habituation for Partitions effect. For each effect that crossed the cluster forming threshold, the following values are presented: the significance probability (p-value, FWE-corrected); the electrode where the effect is most sustained through time; the time of the peak for the most sustained electrode; the t-value at that peak; and the summed t-values of the significant cluster. Clusters are ordered (from left to right) from most to least significant; PC = positive cluster found within the ROI. * indicates p≤0.05.

| *Headache x Habituation for Partitions* |  |  |
| --- | --- | --- |
| *Statistic* | *PC1* | *PC2* |
| *p* | **0.0033\*** | 0.5415 |
| *Most sustained electrode* | A29 | A28 |
| *Time of peak of most sustained* | 133ms | 247ms |
| *Max T(34)-value* | 3.9 | 0.3 |
| *Cluster Stat* | 855 | 13 |

Figure 10(b) shows the scalp topography maps through time for the significant cluster, over a range of 56-256ms after stimulus onset. As seen in both figures HeadacheHabitExploratory(a) and HeadacheHabitExploratory(b), the effect starts at 110ms and lasts until 185ms, sitting over the occipital lobe, with the “peak” electrode being A29.

**Figure 10:**
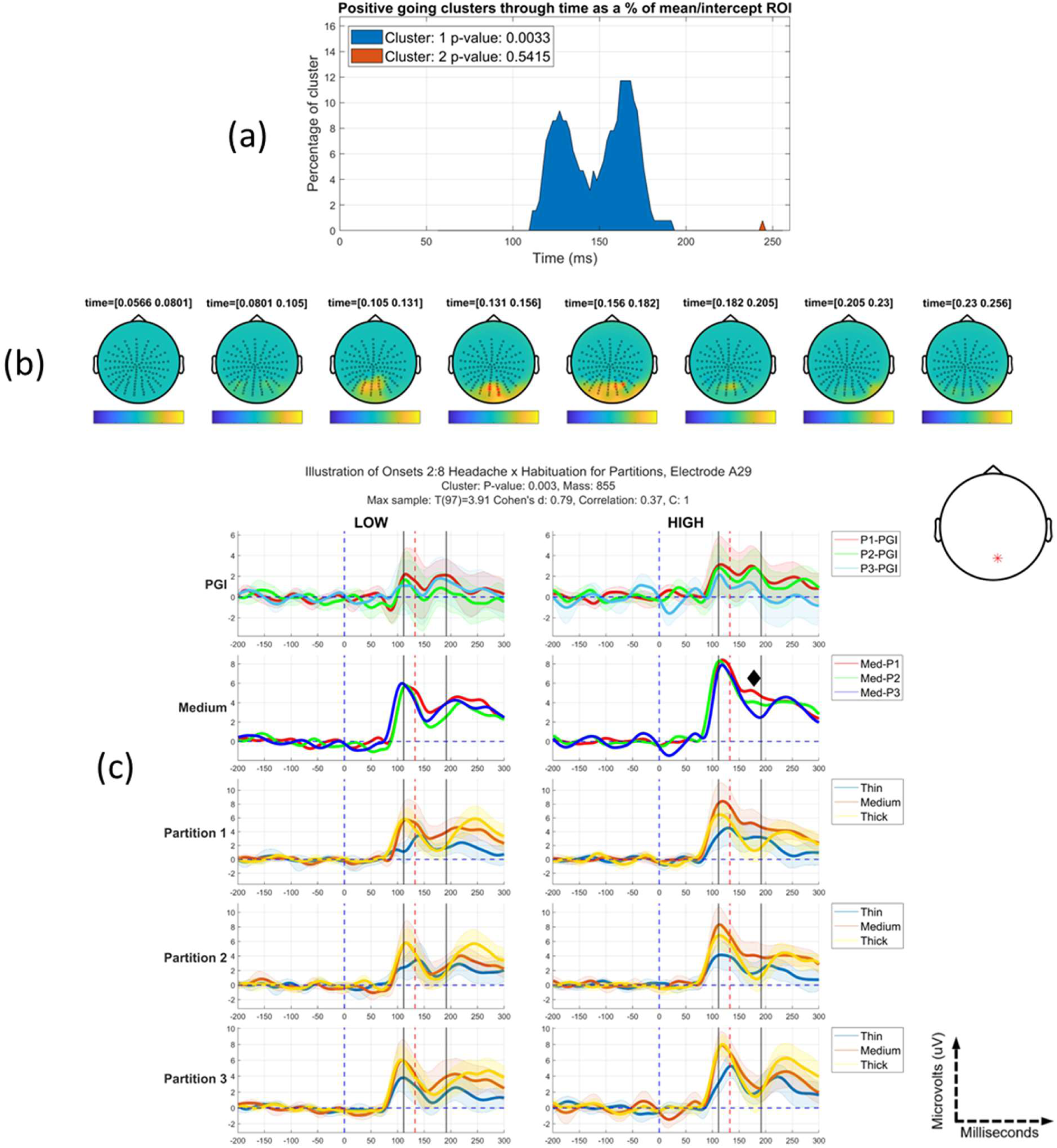
(Unorthogonalized) Headache x Habituation for Partitions effect: (a) Positive going clusters through time as a proportion of the mean/intercept region-of-interest. (b) Topographic maps through bounding time window, 56-256ms after stimulus onset, with time step between each map approximately 25ms. A positive-going cluster is observed at posterior electrodes, specifically in the region of 110-185ms. (c) Grand average time-series of the PGI and individual conditions at the electrode that is the most continuously significant through time. A median split was taken and plots on the left side are for those low on the headache factor and the right side for those high on the factor. The shaded regions represent 95% confidence intervals for each time-series. The black lines indicate the start and end of the most significant cluster through time, with the red dashed line indicating the peak of the effect. Partitions are indicated by P1, P2 and P3. Med_Pi indicates the medium stimulus for partition i. C: j indicates the cluster number. In panel (c), during the period of the significant cluster (approx. 110ms to 185ms) and after it, the High Group (right side of panel (c)) show a habituation pattern that is not present for the Low Group (left side of panel). This is most easily seen in the top row of panel (c), although the biggest reduction in amplitude is from the second to the third partition, suggesting a later quenching of hyper-excitation during the experiment. The second row of panel (c), where medium is plotted alone, suggests the same pattern, but not for the entire period of significance. This suggests that changes in Thick and Thin contribute at some periods during the analysis segment to the significance of the cluster.

Figure 10(c) displays the grand average time-series for the most continuously significant electrode in the cluster, divided into high and low according to a median split on participant scores on the Headache factor. The first two rows of Figure 10(c) show that we are observing a pattern similar to our characteristic habituation pattern. That is, for the low group, the differences between partitions are small and fail to exhibit a consistent change through the partitions (e.g., with the middle partition, 2, the lowest). In contrast, partition one is elevated for the high group. However, while we often see full habituation by partition 2, here there is evidence that the high group does not fully habituate until the 3rd partition. Additionally, the high group in the 2nd row shows that the habituation pattern for the Medium stimulus does not last across the full window of significance, a phenomenon that suggests some involvement of Thick and Thin in the change in PGI through the partitions. The second half of the window of significance does however show the habituation pattern we are interested in for the Medium stimulus (see diamond marker). Thus, these time series suggest hyperexcitation that is specific to the high headache group, which habituates through the course of the experiment. Additionally, the effect on headache reported in Tempesta et. al. (2021) occurs within the time-space region of the effect presented here, suggesting that what we are identifying here is in some respects a decomposition of the earlier effect across partitions. Finally, since this effect is over an unorthogonalized Headache x Habituation regressor, this effect should be considered exploratory.

#### 3.3.2. Headache by Sensitization for Onsets interaction regression

Similarly, to the Headache x Habituation over Partitions effects, we present Headache by Sensitization for Onsets interaction regression results.

##### 3.3.2.1. Orthogonalized Headache x Sensitization for Onsets

One significant positive-going cluster was identified for the orthogonalized Headache x Sensitization for Onsets regression (see Figure 11) over occipital lobe, within the mean/intercept ROI. This cluster spanned 125-205ms. The (FWE-corrected) *p-*value is 0.015, with the peak of the effect found at 174ms and a *t(103)*-value of 3.61 (see Table 5).

**Figure 11:**
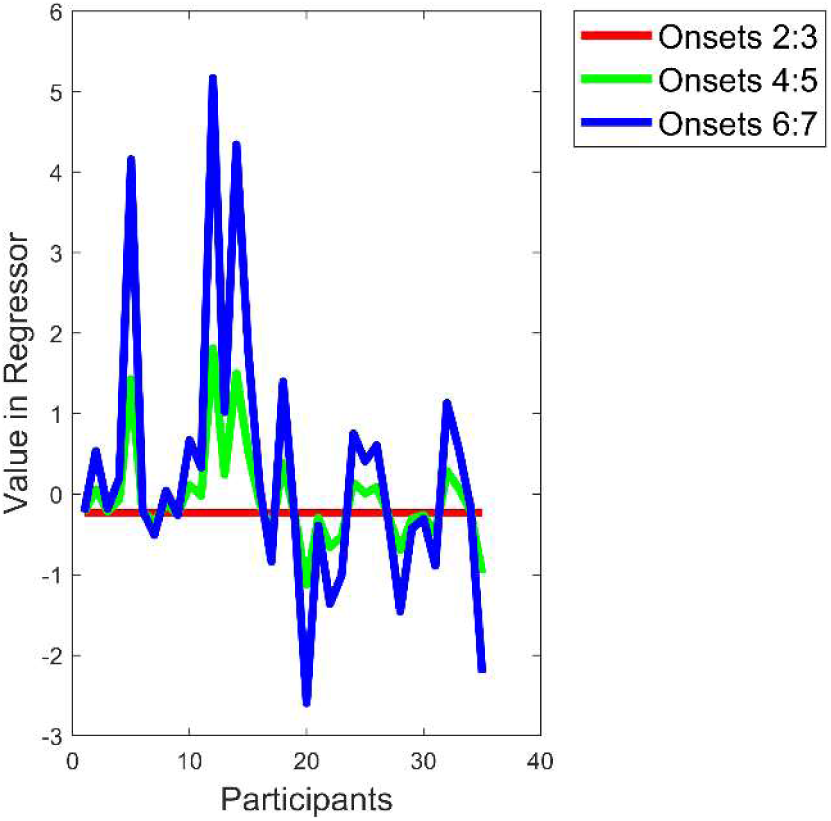
Orthogonalized Headache x Sensitization for Onsets regressor. The x-axis represents the participants and the y-axis the design matrix value assigned to them for each Onset-pair.

**Table 5.** Collective results for the positive (+1) tail of the Headache x Sensitization for Onsets effect. For each effect that crossed the cluster-forming threshold, the following values are presented: the significance probability (p-value, FWE-corrected); the electrode where the effect is most sustained through time; the time of the peak for the most sustained electrode; the t-value at that peak; and the summed t-values of the significant cluster. Clusters are ordered (from left to right) from most to least significant. PC = positive cluster found within the ROI. * indicates p≤0.05.

| *Headache x Sensitization for Onsets* |  |  |
| --- | --- | --- |
| <u>Statistic</u> | <u>PC1</u> | <u>PC2</u> |
| *p* | **0.015\*** | 0.1276 |
| *Most sustained electrode* | A28 | B6 |
| *Time of peak of most sustained* | 174ms | 90ms |
| *Max T(34)-value* | 3.6 | 3.2 |
| *Cluster Stat* | 521 | 113 |

As seen in both figures OrthogHeadacheOnsets(a) and OrthogHeadacheOnsets(b), the effect of interest starts at 125ms and lasts until 205ms, sitting over occipital lobe. Figure 12(c) displays the grand average time-series for the most continuously significant electrode in the cluster (A28), divided into high (right column) and low (left column) according to a median split on the participant scores for the Headache factor.

**Figure 12:**
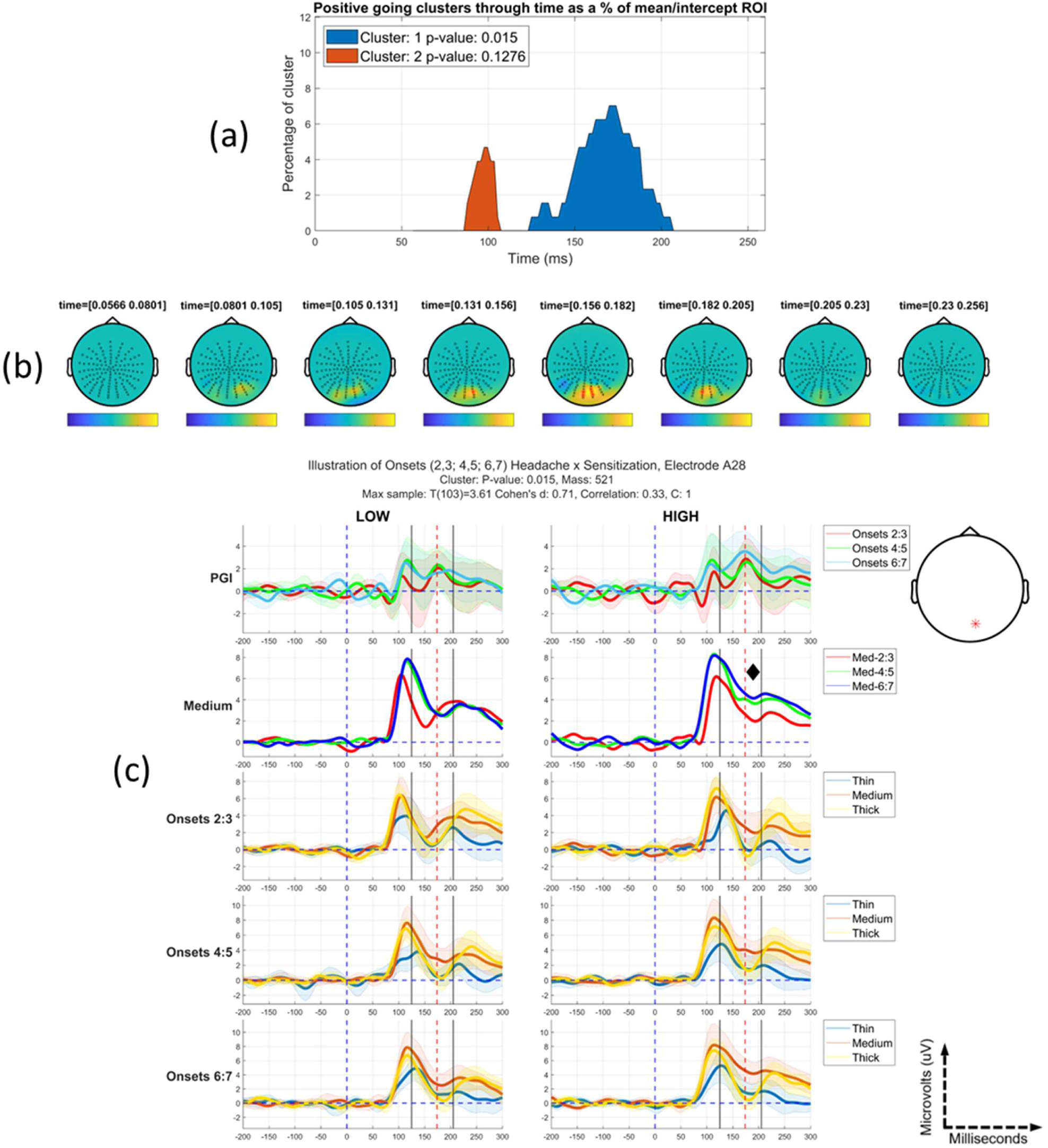
Orthogonalized Headache x Sensitization for Onsets effect: (a) Positive-going clusters through time, with each as a proportion of the mean/intercept region-of-interest. (b) Topographic maps through bounding time window, 56-256ms after stimulus onset, with time step between each map of approximately 25ms. A substantial positive-going cluster is observed at posterior electrodes, which is large between 125 – 205ms. (c) Grand average time-series of the PGI and individual conditions at the electrode that is the most continuously significant through time. The shaded regions represent 95% confidence intervals for each time-series. A median split was taken and plots on the left side are for those low on the headache factor and the right side for those high on the factor. The black lines indicate the start and end of the most continuously significant cluster through time, with the red dashed line indicating the peak of the effect. Med_ij indicates the medium stimulus for onsets i and j. C: k indicates the cluster number. The first row of panel (c) indicates some element of sensitisation for both low and high groups. This is most clearly observed as an elevated PGI for the last onset-pair (6:7). The second row of panel (c), shows that this elevation is also observed for the medium stimulus, making these clean demonstrations of hyper-excitation. However, the hyper-excitation pattern tends to be early and brief for the low group, while it is more sustained and also present over later time periods for the high group.

The 1^st^ and 2^nd^ rows of Figure 12(c), show a compelling differential sensitization effect. This is particularly the case, if one focuses on the region of significance from about 150ms (see diamond marker) onwards (indeed, the substantive part of cluster one in Figure 12(a) does not start until approaching 150ms). For the high group, the response to both PGI (first row) and medium (second row) generally increases through Onset-pairs (from 2:3 to 4:5 to 6:7), while from soon after 150ms, for the low group, there is no clearly evident sustained change through Onset-pairs. This finding suggests a tendency (from 150ms) for hyper-excitation to emerge through repeated presentation of an aggravating stimulus only for those susceptible to headaches. This effect could be considered a decomposition of the headache factor effect in (Tempesta, Miller, Litvak, Bowman, & Schofield, 2021)according to change through Onsets, from second onwards.

##### 3.3.2.2. Non-orthogonalized Headache x Sensitization for Onsets (Exploratory)

One significant positive-going cluster was identified for the Headache x Sensitization for Onsets regression (see Figure 13) over occipital lobe within the mean/intercept ROI (see Figure 14). This effect spanned 125-215ms. The (FWE-corrected) *p-*value is 0.0057, with the peak of the effect found at 142ms with a *t(103)*-value of 4.3 (Table 6).

**Figure 13:**
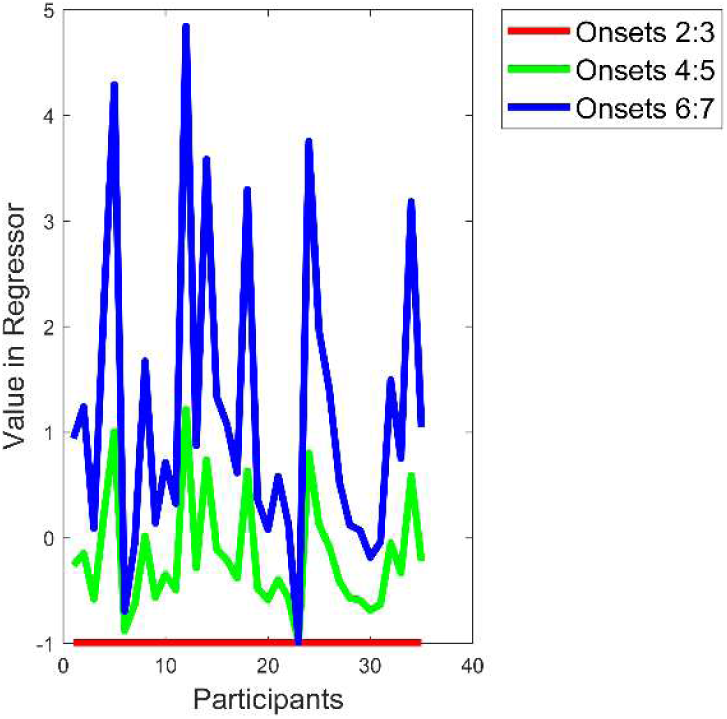
Headache x Sensitization for Onsets regressor. The x-axis represents the participants and the y-axis the design matrix value assigned to them for each Onset.

**Figure 14:**
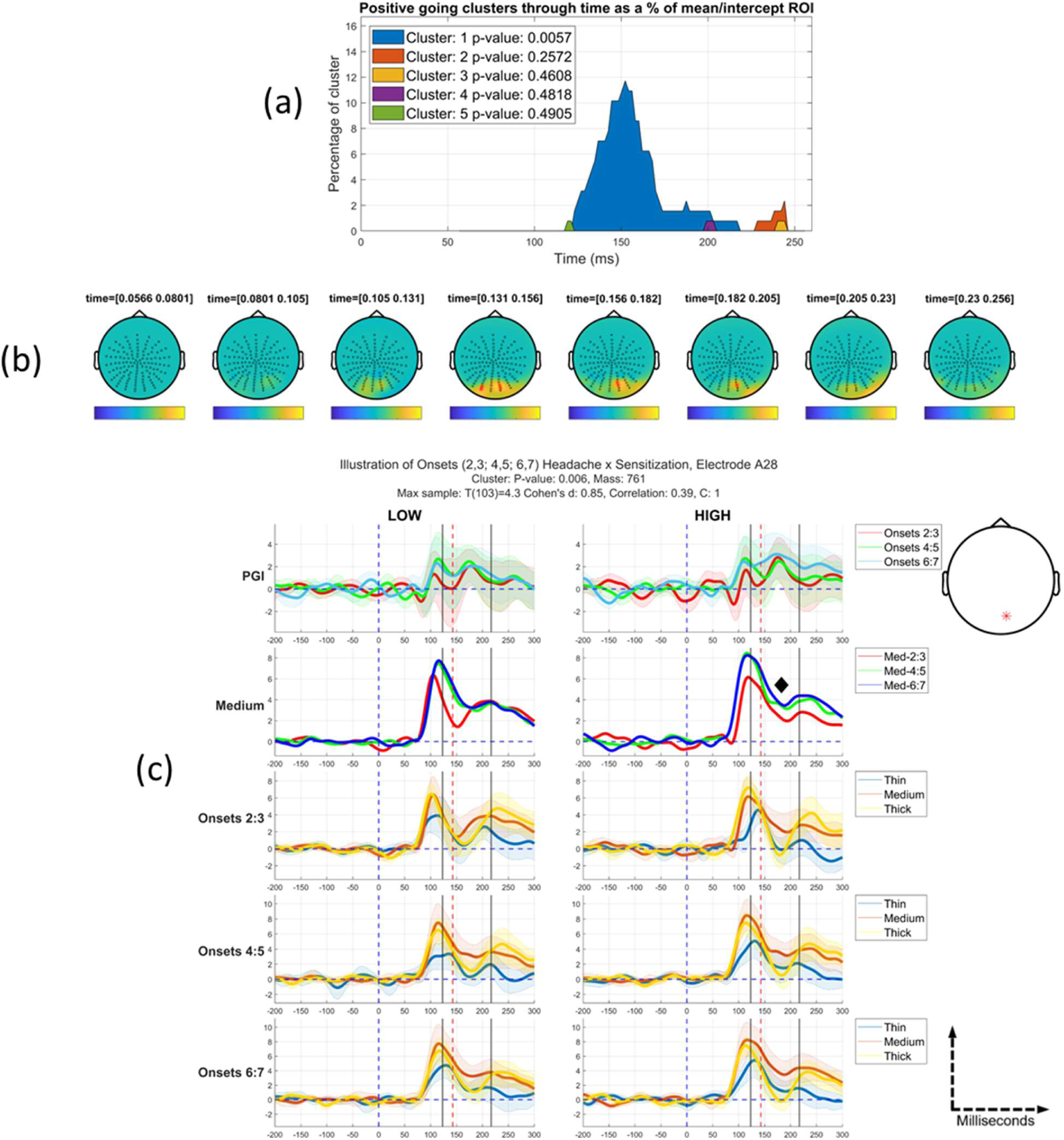
Headache x Sensitization for Onsets effect: (a) Positive-going clusters through time, with each as a proportion of the mean/intercept region-of-interest. (b) Topographic maps through bounding time window: 56-256ms after stimulus onset, with time step between each map approximately 25ms. A positive-going cluster is observed at posterior electrodes. (c) Grand average time-series of the PGI and individual conditions at the electrode that is the most continuously significant through time. The shaded regions represent 95% confidence intervals for each time-series. The black lines indicate the start and end of the most significant cluster through time, with the red dashed line indicating the peak of the effect. Med_ij indicates the medium stimulus for onsets i and j. C: k indicates the cluster number. The first row of panel (c) indicates some elements of sensitisation for both low and high groups. This is most clearly observed as an elevated PGI for the last onset-pair (6:7). The second row of panel (c), shows that this elevation is also observed for the medium stimulus, making these clear demonstrations of hyper-excitation. However, the hyper-excitation pattern tends to be early and brief for the low group, while it is more sustained and over later time periods for the high group. (Note, orthogonalization has a small impact on the median split, since the scores associated with participants changes. Consequently, the time-series in this figure are not exactly the same as those in Figure 12.)

**Table 6.** Collective results for the positive (+1) tail of the Headache x Sensitization for Onsets effect. For each effect that crossed the cluster forming threshold, the following values are presented: the significance probability (p-value, FWE-corrected); the electrode where the effect is most sustained through time; the time of the peak for the most sustained electrode; the t-value at that peak; and the summed t-values of the significant cluster. Clusters are ordered (from left to right) from most to least significant; PC = positive cluster found within the ROI. * indicates p≤0.05.

| *Headache x Sensitization for Onsets* |  |  |  |  |  |
| --- | --- | --- | --- | --- | --- |
| <u>Statistic</u> | <u>PC1</u> | <u>PC2</u> | <u>PC3</u> | <u>PC4</u> | <u>PC5</u> |
| *p* | **0.0057\*** | 0.2572 | 0.4608 | 0.4818 | 0.4905 |
| *Most sustained electrode* | A28 | A28 | D32 | B9 | A13 |
| *Time of peak of most sustained* | 142ms | 244ms | 244ms | 200ms | 121ms |
| *Max T(34)-value* | 4.3 | 3.1 | 2.6 | 2.3 | 2.5 |
| *Cluster Stat* | 761 | 37 | 8 | 7 | 5 |

As seen in Figure 14(a) and HeadacheOnsetsExploratory(b), the effect of interest starts at 125ms lasting until 215ms and sitting over the occipital lobe with the peak electrode being A28. Figure 14(c) displays the grand average time-series for the most continuously significant electrode in the cluster, divided into high and low according to a median split on the participant scores on the Headache factor.

We again see a differential sensitization pattern, which looks convincing for the Medium stimulus (i.e., second row of Figure 14 (c)) from about 155ms onwards (see diamond marker), with only the high group showing strong sensitization. However, the pattern for PGI (i.e., first row of Figure 14 (c)) is not quite so convincing, with the high group only intermittently exhibiting clear sensitization across the partitions. Thus, when the medium stimulus is placed in comparison with its control stimuli (thick and thin) the effect is not so clear cut. (Note, this unorthogonalized version of this interaction regressor, has a change in mean amplitude across partitions built into it. Consequently, even if the headache pattern does not fit well, an effect can be carried by simple change in mean amplitude across partitions. This might particularly be what is driving the effect at the point where it is biggest (red dashed vertical line). We consider this issue further in the Discussion.)

### 3.4 Three-way Interaction: Unorthogonalized Headache x Habituation (for Partitions) x Sensitization (for Onsets) [exploratory]

The Headache x Habituation (for Partitions) x Sensitization (for Onsets) interaction was conducted specifically to see whether we could connect the two-way interactions that we have observed: Headache x Habituation (for Partitions) and Headache x Sensitization (for Onsets). Could we find evidence that the sensitization effect we observed for Onsets, habituates across the course of the experiment (i.e., across Partitions)? We did not test any other three-way interactions. Only the unorthogonalized three-way interaction came out as significant. (The non-significant orthogonalized version of this contrast is presented in Appendix 5).

One significant-positive going cluster was identified for the Headache x Habituation (for Partitions) x Sensitization (for Onsets) effect; see regressor in Figure 15. The effect sits over the occipital lobe within the mean/intercept ROI. The effect for the most positive going cluster spanned 125-195ms. The (FWE-corrected) *p*-value is 0.0133, with the peak of the effect found at 176ms, with a *t(232)*-value of 3.4 (see Table 7).

**Figure 15:**
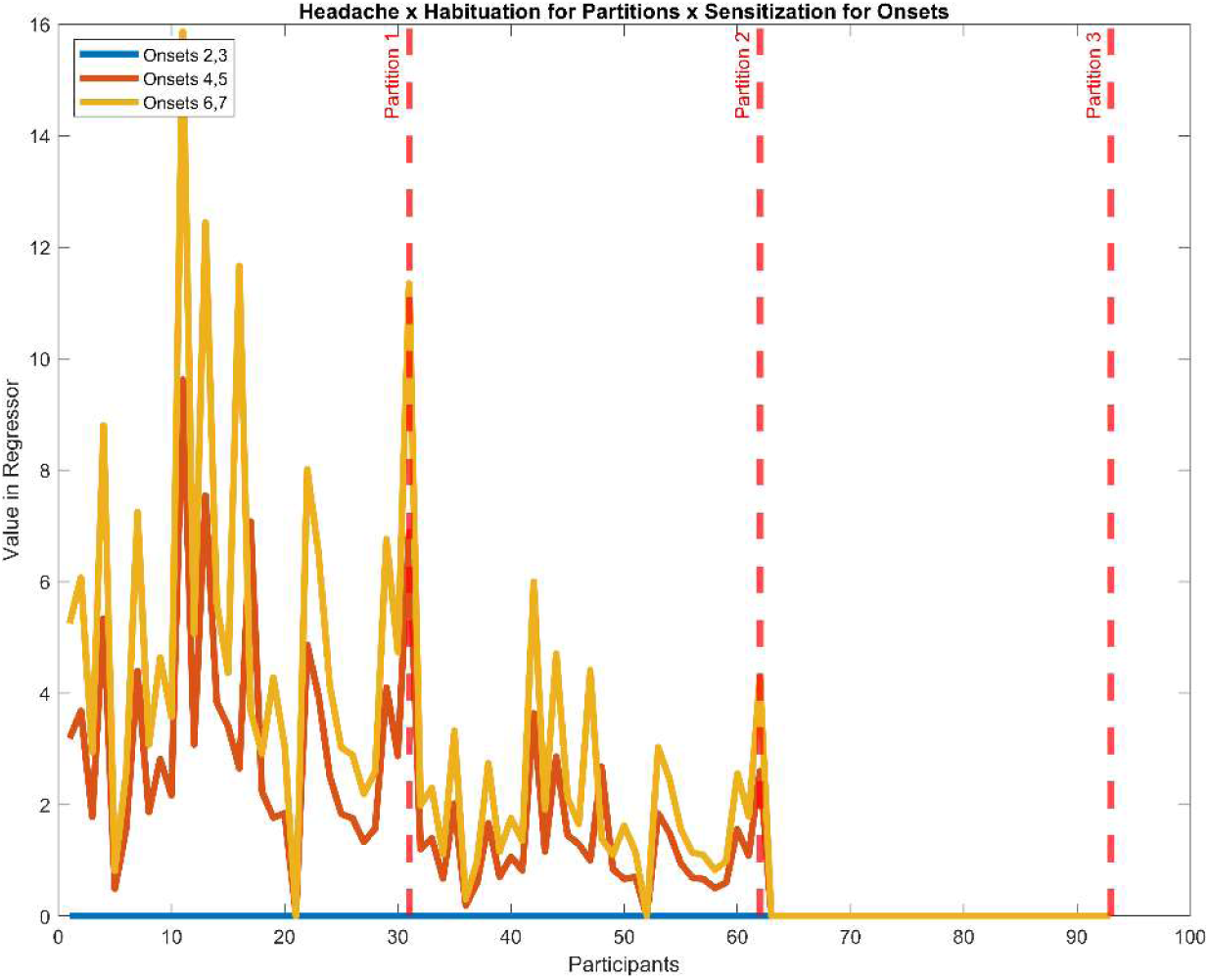
(Unorthogonalized) Headache x Habituation (for Partitions) x Sensitization (for Onsets) effect: (a) The x-axis represents participants, all of which appear three times, once for each partition. The y-axis represents the design matrix value assigned to each participant for each onset in each partition. Onsets are indicated by the blue, orange, and yellow lines (2:3; 4:5; 6:7) with partitions segmented by the red dashed lines. Note, in the final partition (number 3), the red and blue lines are “under” the yellow line.

**Table 7.** Collective results for the positive (+1) tail of the Headache x Habituation for Partitions x Sensitization for Onsets effect. For each effect that crossed the cluster forming threshold, the following values are presented: the significance probability (p-value, FWE-corrected); the electrode where the effect is most sustained through time; the time of the peak for the most sustained electrode; the t-value at that peak; and the summed t-values of the significant cluster. Clusters are ordered from most to least significant; PC = positive cluster. * indicates p≤0.05.

| *Headache x Habituation for Partitions x Sensitization for Onsets* |  |  |  |  |
| --- | --- | --- | --- | --- |
| *Statistic* | *PC1* | *PC2* | *PC3* | *PC4* |
| *p* | **0.0133\*** | 0.646 | 0.4678 | 0.4688 |
| *Most sustained electrode* | A29 | A29 | B10 | A10 |
| *Time of peak of most sustained* | 176ms | 132ms | 230ms | 104ms |
| *Max T(34)-value* | 3.4 | 3.4 | 2.3 | 2.3 |
| *Cluster Stat* | 542 | 179 | 11 | 8 |

Figure 16 (b) shows the scalp maps through time during our bounding window (56-256ms after stimulus onset) for the significant cluster. As seen in both figures *HeadacheThreeWayExploratory* (a) and *HeadacheThreeWayExploratory* (b), the effect starts at 125ms lasting until 195ms, positioned over the occipital lobe with the peak electrode being A29.

**Figure 16:**
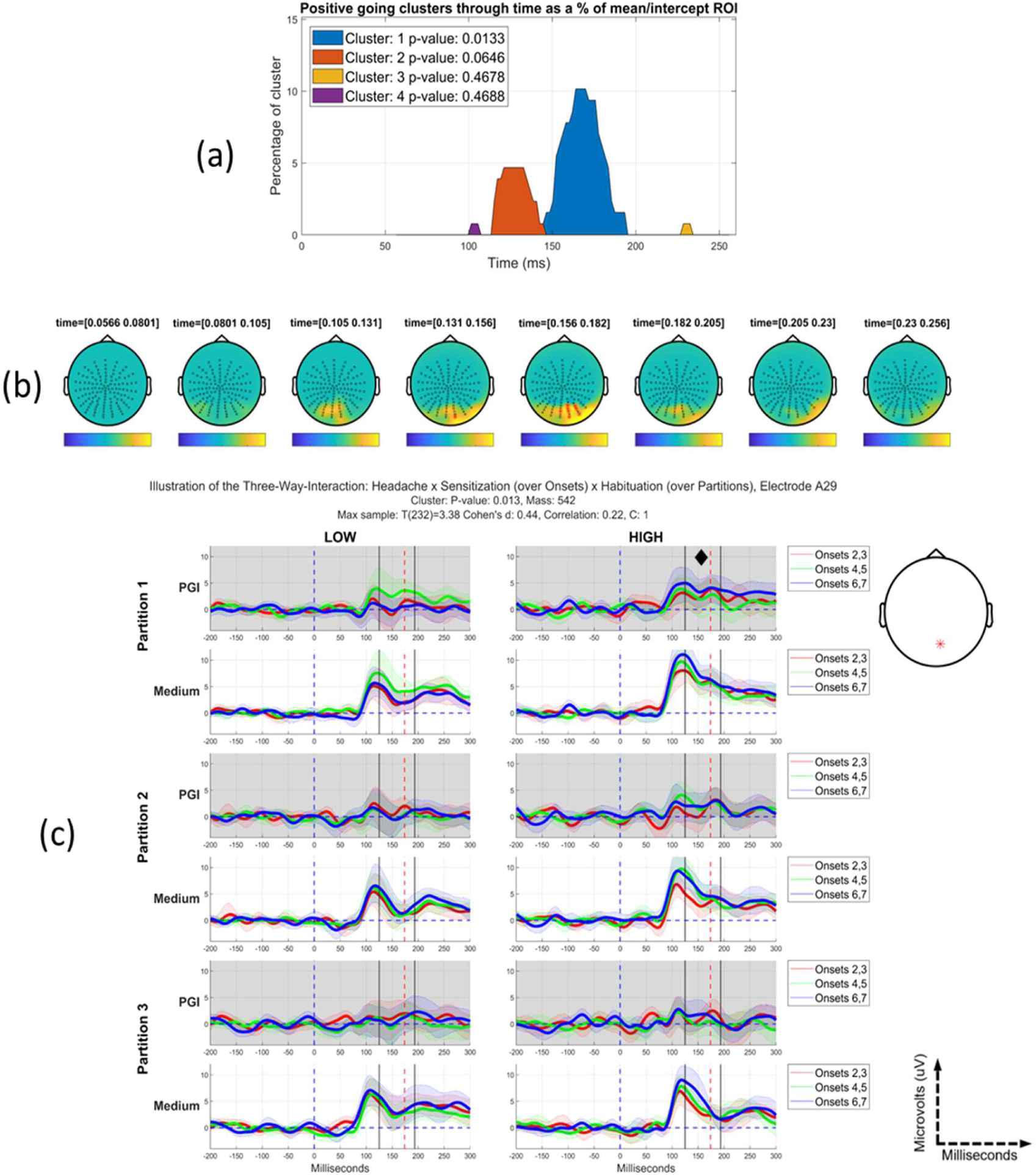
Headache x Habituation (for Partitions) x Sensitization (for Onsets) effect: (a) Positive going clusters through time as a proportion of the mean/intercept region-of-interest. (b) Topographic maps through bounding time window, 56-256ms after stimulus onset, with time step between each map approximately 25ms. A positive-going cluster is observed at posterior electrodes, specifically in the region of 125-195ms. (c) Grand average time-series of the PGI and Medium stimulus at the electrode that is most continuously significant through time. The shaded regions represent 95% confidence intervals for each time-series. A median split was taken and plots on the left side are for those low on the headache factor and the right side for those high on it. The black vertical lines indicate the start and end of the most significant cluster through time, with the red dashed line indicating the peak of the effect. Rows showing the PGI are all in grey; those showing the Medium stimulus are in white. Partition 1 is indicated by the first two rows, partition 2 the second two and partition 3 the last two. Within each partition, the PGI and medium stimulus is highlighted for onsets 2:3, 4:5 and 6:7. The most direct way to understand the effect observed, is to concentrate on the PGI rows (presented in grey) in panel (c): the first (partition 1), third (partition 2) and fifth (partition 3) rows. Although certainly not a completely “consistent” pattern through time, in the time-period of the significant cluster and also before and after it, there is some evidence of partitions differentially (across high and low groups) modulating the effect on onsets. For example, we do see a pretty sustained sensitization effect across onsets in the top right plot in panel (c) (i.e., High group, partition 1). In that plot, onsets-pair 6:7 (blue line) exhibits an elevated PGI throughout the period of the EEG transients, apart, though, from at the point of maximum effect, the red dashed line; we return to this issue in the Discussion. Additionally, onsets-pair 6:7 (blue line) is in no way elevated in the top left plot (i.e., Low group, partition 1), and never for a sustained period in either group for partitions 2 or 3.

Figure 16 (c) displays the grand average time-series for the most continuously significant electrode in the cluster, segmented into high (right-side) and low (left-side) according to a median split on the participant scores for the Headache factor. The most direct way to understand the effect is to concentrate on the PGI rows (shown in grey) in panel (c): the first (partition 1), third (partition 2) and fifth (partition 3) rows. Although certainly not a *completely* “consistent” pattern through time, in the time-period of the significant cluster and also before and after it, there is evidence of partitions differentially (across high and low groups) modulating the effect on onsets. For example, we do see a pretty sustained sensitization effect across onsets in the top right plot in panel (c) (High group, partition 1, see diamond marker). In that plot, onset-pair 6:7 (blue line) exhibits an elevated PGI throughout the period of the EEG transients, apart, though, from at the point of maximum effect, the red dashed line; we return to this issue in the Discussion. Additionally, onsets-pair 6:7 (blue line) is in no way elevated in the top left plot (i.e., Low group, partition 1), and never for a sustained period in either group for partitions 2 (row 3) and 3 (row 5). This pattern of responses is weaker for the Medium stimulus, when considered on its own; see rows 2, 4 and 6 (white background rows). This raises the possibility that the three-way interaction on PGI is at least partially caused by changes in Thick and Thin, rather than just on Medium.

Additionally, the second cluster observed in Figure 16(a), which exhibits a trend towards significant (p-value=0.0646), is largest at the same electrode as the first cluster (A29) (see Table 7), but somewhat earlier in time (between approx. 115ms and 150ms). Thus, this second cluster can also be observed in Figure 16(c), providing evidence that the three-way interaction effect we are observing is extended in time.

Although not a perfect match, the effect we present in Figure 16 exhibits some overlap with the space and time of the Headache x Sensitization for Onsets effect, whether orthogonalized or not (see figures *OrthogHeadacheOnsets* and *HeadacheOnsetsExploratory*), and the Headache x Habituation for Partitions effect when not orthogonalized (see Figure 10). This suggests that the three-way interaction we observe in Figure 16 is part of the same phenomenon that generates the two two-way interactions.

This effect raises the possibility that (differentially) for headache sufferers, hyper-excitation increases through repetition of onsets when they first start viewing pattern-glare stimuli. However, the brains of these participants habituate, leading to reduced hyper-excitation, over a longer time period.

### 3.5 Summary of Contrasts

Table 8 summarises the temporal effects we observed and failed to observe. In our results section, we have focussed on sensitization through the onsets and habituation through the partitions. However, we did also explore the opposite patterns: habituation through the onsets and sensitization through the partitions, and importantly, as shown in Table 8, no effects (even just crossing the first level threshold) were found for these alternative patterns of change through time. This adds credence to our position that the dominant pattern in our data is short-term sensitization (through onsets), accompanied by longer-term habituation (through partitions).

**Table 8.** Summary of statistically significant clusters across both time granularities (onsets and partitions) and directionality (habituation and sensitization). The Pure Time contrasts exhibit a change through time, without crossing with the Headache factor; see Figure 4; thus, they could be considered as Main Effects of the analysis. (The full details of the Pure Time contrast are presented in a companion paper (Dogan, et al., 2023).) “No cluster found” indicates no clusters formed, i.e., no samples crossed the first level (cluster-forming) threshold. “Non-significant cluster” indicates that there were samples (time-space points) that crossed the cluster-forming threshold, but the resulting clusters did not survive second-level family-wise error correction.

|  | Partitions |  | Onsets |  |
| --- | --- | --- | --- | --- |
| Contrast | Habituation | Sensitization | Habituation | Sensitization |
| **Pure Time Effect** | Non-significant cluster (see results) | No cluster found | No cluster found | No cluster found |
| **Headache x Time (unorthogonalized)** | Significant Cluster Found (see results) | No cluster found | No cluster found | Significant Cluster Found (see results) |
| **Headache x Time (orthogonalized)** | Non-significant cluster (see appendix) | No cluster found | No cluster found | Significant Cluster Found (see results) |

## 4. Discussion

We based our analyses on four main research questions about the electrophysiological correlates of pattern-glare in relation to headache susceptibility. We discuss how each of these questions stands in the light of our findings.

### 4.1. Mean/Intercept and Headache (main) effects (question/contrast 1)

#### Mean/Intercept

As shown in Figure 6, across the whole group of participants, we see a sustained highly-significant “hyper-excitation” effect. Medium is above the average of Thick and Thin throughout this single cluster, suggesting that the effect is driven by an elevated response to Medium, consistent with prior research (Newman Wright, Wilkins, & Zoukos, 2007; Wilkins A., Intermittent illumination from visual display, 1986; Schoenen, Wang, Albert, & Delwaide, 1995; Gerber & Kropp, 1995; Tempesta, Miller, Litvak, Bowman, & Schofield, 2021). The effect is most potent (at some points, above 20% of entire volume) in the occipital lobe, centred at A26, lasting effectively from the start of the onset transients (∼100ms) to the end of the analysis segment (256ms) and, in fact, beyond segment end; see (Jefferis, 2023). There are two effect peaks, one around 120ms (coinciding with the P1) and the other around 180ms (coinciding with the following N1); see Figure 6(a).

This finding may relate to previous MEG (Adjamian, et al., 2004) findings. Adjamian et al. (2004) detected (broadband gamma) oscillatory correlates of hyper-excitation over the visual cortex of a healthy group of participants, with gratings close to the spatial frequency of our Medium stimulus eliciting the highest increase in power. Further work should attempt to relate our time domain and Adjamian et al’s frequency domain effects. For example, do the amplitudes of these two effects correlate across participants?

For our findings, a potential confound that it is important to exclude is the possibility that the evoked response effects that we observe are due to eye movements. A detailed analysis on the eye electrodes (VEOG/HEOG) is included in Appendix 1 “Ruling out eye confounds”.

#### Factor main effect

We also observed an effect directly on the Headache factor; see Figure 8, which is significant, but considerably weaker than the Mean/Intercept effect. This sensitivity to Headache susceptibility is found during the second peak of the Mean/Intercept, around 180ms after stimulus onset, during the first negativity. This effect suggests increased hyper-excitation (i.e., pattern-glare index) for those more susceptible to have headaches. This finding is broadly consistent with previous findings that migraineurs exhibit higher amplitude early visual-evoked potentials (Oelkers, et al., 1999).As discussed in more detail in (Tempesta, Miller, Litvak, Bowman, & Schofield, 2021), two studies have previously measured ERPs in a headache prone population using pattern-glare-like stimuli. ( (Fong, Law, Braithwaite, & Mazaheri, 2020) found differences between migraine sufferers and controls at around 200- and 400ms post stimulus onset. Their migraine group showed greater negativity at 200ms for high-frequency gratings (13 c/deg). Indeed, their main findings were on the high-frequency grating, while in contrast, the findings we report here occur with the clinically relevant, medium-frequency grating (3 c/deg). However, as elaborated in (Tempesta, Miller, Litvak, Bowman, & Schofield, 2021), it is difficult to compare our effects to Fong et al’s. This is because Fong et al did not use repeated onsets and so could not observe an analogue of the effects reported here, which are across Onsets 2 to 8, during which time, unlike our Onset 1 and Fong et al’s study, the stimulus presented is completely predictable.

Furthermore, (Haigh, Chamanzar, Grover, & Behrmann, 2019) found larger amplitude N1 and N2 ERP components in a migraine group viewing chromatic gratings. These enlarged responses were associated with higher discomfort ratings when viewing the stimuli.

#### Comparison to Tempesta et al

The mean/ intercept reported here corresponds to the mean/ intercept effect observed in (Tempesta, Miller, Litvak, Bowman, & Schofield, 2021); and there is no surprise that we observe this again, since we are analysing the same data. However, the effect here is larger, because Fieldtrip enables a more appropriate cluster inference for EEG data, where a first-level (cluster-forming) threshold can be set at 0.025, rather than the 0.001 required in SPM (Flandin & Friston, 2017). The signal in scalp-EEG tends to be relatively low amplitude, but broad in space and time (i.e., smooth). SPM’s 0.001 first level threshold tends to be too strict to detect such low amplitude-broad effects.

Additionally, our analysis on the (pure) headache factor (a main-effect in the design) also repeats an analysis presented in (Tempesta, Miller, Litvak, Bowman, & Schofield, 2021), with the same data. This headache factor effect was significant in Tempesta et al (t(30) = 3.34, p = 0.047, FWE cluster-level corrected, with Random Field Theory and small volume correction), and similarly so in this paper (p = 0.046, see details in Table 3).

However, all the interaction effects that we report in this paper are new analyses. Thus, the key original contribution of this paper is the identification of profiles of change through time, i.e., habituation and sensitization, and how those profiles interact with Headache susceptibility.

One aspect of the (Tempesta, Miller, Litvak, Bowman, & Schofield, 2021) findings was the demonstration of increased temporal jitter for the Medium stimulus for those high on the headache factor. It is a possibility that the interaction effects we observe reflect changes in temporal jitter over fine (across onset-pairs) and coarse (across partitions) timeframes for those high on the Headache factor. However, confirming this is challenging, since calculation of phase coherence is impacted by changes in power, and we do indeed observe such changes through the course of our experiment.

### 4.2. Headache by Habituation for partitions interaction effect (question/contrast 2)

Our second finding suggests that this hyper-excitation over the visual cortex decreases (i.e., habituates) through the course of the entire experiment; see Figure 10. Furthermore, this decrease is modulated by headache sensitivity, with those high on the Headache factor habituating more. This effect suggests brain adaptation on a relatively macroscopic temporal frame, as it was demonstrated over the experiment’s blocks that lasted for 10-12 minutes each.

This finding demonstrates the brain’s ability to habituate to an aggravating visual stimulus. Evidence of habituation occurring in the visual cortex already exists (Obrig, et al., 2002; Schoenen, Wang, Albert, & Delwaide, 1995)) and in some cases, the presence of habituation was also discovered for migraine sufferers (Khalil, 2000; Omland, et al., 2016). However, a conflict exists between studies, as previous work has suggested that migraineurs do not habituate (Adjamian et al., 2004; Schoenen et al., 1995). We shortly return to the question of why we observe the particular pattern of change through time that we do; see subsection 4.6. “Potential Explanation of Findings”.

Finally, it is important to also acknowledge that this Headache-by-Habituation for partitions effect is only significant unorthogonalized. Consequently, this is an effect where replication is certainly required.

### 4.3. Headache by sensitization for onsets interaction effect (question/contrast 3)

As shown in Figures OrthogHeadacheOnsets and HeadacheOnsetsExploratory, we have obtained evidence for a sensitization effect arising from stimulus repetition that is differentially present for participants susceptible to headaches. This suggests that repeated presentation of a pattern-glare stimulus, with repetition onsets spaced by around four seconds, drives hyper-excitation in the brain to increase through these repetitions for headache sufferers.

(Mickleborough, et al., 2014) conducted an experiment in which a series of visually complex images were presented to a population of clinically defined migraineurs and healthy participants. Similar to our findings, participant ERPs for the migraineurs relative to the healthy participants exhibited sensitization quickly after stimulus onset, for the duration of the experiment. This sensitization was present for the initial blocks, however, at later stages during the experiment, migraineur’s ERPs started to show a habituation effect. It is important to highlight that, unlike our experiment in which we recruited from the general population and recorded self-report headache susceptibility, the participants employed by Mickleborough et. al. were clinically confirmed migraineurs. Additionally, the experimental paradigm used by Mickleborough et. al. is much shorter in overall duration, which may explain why sensitization is present throughout the entire experiment. Finally, since this effect was significant for both the orthogonalized and unorthogonalized contrasts, with identified-clusters that strongly overlap in space and time, we have to consider this effect more reliable than the Headache-by-Habituation for partitions effect reported in the previous subsection.

### 4.4. Headache-by-habituation for partitions by-sensitization for onsets interaction effect (question/contrast 4)

Because both the Headache-by-habituation for partitions and the Headache-by-sensitization for onsets interactions were significant, with similar scalp topographies and overlap in time, a natural next question is whether these two two-way interactions interact with one another. To explore this, we specifically looked at the Headache-by-habituation for partitions by-sensitization for onsets interaction effect.

Consequently, we have found evidence for the possibility that (differentially) for headache sufferers, hyper-excitation increases through repetition of onsets when they first start viewing pattern-glare stimuli. However, the brains of these participants habituate, leading to reduced hyper-excitation, over a longer time period. This raises the possibility that those high on the Headache factor are aggravated, i.e., hyper-excited, by repeated presentation when they start the experiment, but that aggravation through repetition dissipates for the rest of the experiment.

Finally, it is important to also acknowledge that the underlying Headache-by-Habituation for partitions effect is only significant unorthogonalized. Additionally, the three-way interaction effect was not strongly evident for the Medium stimulus when visualised on its own (see Figure 16, specifically the white backgrounds in panel (c)), suggesting that changes in Thick and Thin also contribute to this three-way interaction on PGI.

### 4.5 Formulation of Interaction Regressors

There are subtleties to our interaction regressors. This is because they involve two aspects, i.e., for those higher on headache, there should be, (i) more *change through time* (i.e., habituation or sensitization for Partitions or Onsets); and (ii) *larger amplitudes*.

Importantly, because our interest is in hyper-excitation, one does need the second aspect, as well as the first. For example, for Headache by Habituation over Partitions, we do not just want to know that those higher on Headache show more of a reducing pattern across partitions (which might go from positive to negative), we also want to know that this reflects a dissipation of excitation *towards zero*, i.e., that the habituation is from an extreme amplitude down to zero. This is, for example, apparent in panel (a) of Figure 1, where the average value (across time-points) in the regressor is only at baseline (i.e., -1) for the lowest participant on Headache (see participant 22, for whom, red, green, blue is all at -1), but this average value increases as one progresses up the Headache regressor.

Although the three-way is more complex, involving two patterns of change through time, these two aspects, change through time and larger amplitudes, are also both inherent to this three-way pattern.

However, a consequence of these “two-aspect” regressors is that effects that are particularly strong in just one of these two aspects can come out significant in the absence of the other. For example, in Figure 16(c) at the point of maximum effect (red vertical dashed line), the three-way is largely driven by the amplitude effect with little evidence of differential change through time. That is, in the top right plot of panel (c), amplitudes are big (relative to the low group), but there is little apparent sensitization. (Although, at other time-points in the cluster, there is differential sensitization present, i.e., for High [right-side], but not for Low [left-side]).

### 4.6 Potential Explanation of Findings

#### First negativity

All our effects involving the Headache factor (i.e., the Headache factor on its own; Headache-by-habituation for partitions; Headache-by-sensitization for onsets; and Headache-by-habituation for partitions by-sensitization for onsets) are largest at A28 or A29, i.e., effectively the same point on the scalp. Additionally, although there is some variability, with all these effects, there is typically a peak close to 180ms. This suggests that all these Headache effects are really arising from the same features in the data, and potentially the same source(s) in the brain. As further discussed in (Tempesta, Miller, Litvak, Bowman, & Schofield, 2021), this time and space region seems to coincide with a negativity in the raw ERPs, which is particularly evident in the Thick condition, but typically not in the PGI plots, since they represent a contrast of raw ERPs.

This negativity can, for example, be observed in the last three rows panel (c) of Figure 12, where the peak of the effect (see dashed red line) picks out the minimum of this negativity. Furthermore, the effects we observe are often associated with the Medium condition exhibiting a much shallower (or even absent) negativity or a change in this negativity through time. Further work needs to be performed, to understand the brain correlates of this negativity and its relationship to feedforward and/or feedback processing in visual cortex.

#### Change through time

Why, then, do we observe the particular pattern of sensitization and habituation that we have reported? A number of points can be made:

1. sensitization over onsets: perhaps the most expected and also strongest effect we observe is sensitization for those high on the headache factor, for the finer time granularity (i.e., onsets), suggesting, a potential deficit in inhibitory mechanisms to control stimulus-driven excitation in striate cortex.
2. habituation over partitions: a more surprising, but perhaps more intriguing, finding is that hyper-excitation seems to habituate through the course of the experiment. This suggests a successful dampening of hyper-excitation over a long enough period of time for our headache sufferers. However, it is important to remember that our cohort is not clinically selected, and thus may not contain extreme headache sufferers, or only a small number if present. Basically, even for those high on the Headache factor, we may be looking at a relatively well-functioning brain. Consequently, it may not be surprising that their visual systems do successfully habituate. Future work should validate this effect on a clinical group of headache/migraine sufferers. Indeed, it may be that those with extreme sensitivity to pattern-glare stimuli, do not habituate over the course of the experiment or may even sensitise, i.e., with correlates of hyper-excitability increasing through the course of the experiment.
3. Lack of habituation for low on factor: the lack of habituation for those low on the headache factor may simply reflect that they have nothing to habituate, i.e., no hyper-excitation in the first place. That is, this does not necessarily imply that they cannot habituate; it is more that they do not have to.

### 4.7. Robustness of Factor Analysis Decomposition

An important question to consider is the reliability of our factor analysis. We have explored this in depth, by 1) comparing with the only real alternative, what we call a flat average weighting of dimensions, based upon intuition, and 2) a bootstrapping procedure to determine whether there is substantially more uncertainty associated with the loadings arising from the factor analysis than from the flat average. On the basis of these two explorations, we do not see evidence for substantial unreliability arising from the factor analysis. These explorations can be found in Appendix 4 (titled “Justification of factor analysis decomposition”).

### 4.8. Limitations

There are a few limitations to the reported work. As the replies to the questionnaires, as well as the discomfort ratings during the experiment, are self-reported, there is subjectivity in the derived factors, for example, headache regularity. Enlarging the data set and collecting headache diaries would increase the reliability of our findings.

Additionally, as already stressed, there is collinearity between the regressors that result from combining the factor scores and partitions. To ensure that no effect in the data is accounted for twice, the headache-partitions interaction and discomfort-partitions interaction regressors had to be orthogonalized with respect to the visual stress-partitions interaction regressor and then with respect to each other. This choice is justified as visual stress was the strongest factor obtained from the factor analysis and consequently, the other two factors were orthogonalized with respect to it. This limitation deems our non-orthogonalized findings as exploratory.

A number of commentators have rightly highlighted the problem of partial presentation of analyses performed (Bishop, 2020). If many contrasts are tested within an experimental or analysis trajectory, but the results of only a few contrasts, typically those that came out significant, are reported, then it is difficult for the reader to assess the vulnerability to false positives. Accordingly, we want the reader to recognise that the results presented in this paper were part of an analysis trajectory that includes results presented in a companion paper (Dogan, et al., 2023), and in both this paper and (Dogan, et al., 2023), we have presented a table summarising all the contrasts run; see Table 8 in this paper and Table 8 in (Dogan, et al., 2023). Importantly, the general pattern in both papers is to observe habituation (but not sensitization) across partitions and sensitization (but not habituation) across onsets, with these temporal patterns increasing with clinical condition (i.e., as one moves up a factor).

Overall, though, given the novelty of the effects identified, all our interactions need to be replicated before they can be considered robust findings.

### 4.9. Future Research

Increasing the size of the dataset would enable us to investigate quartiles of participants, rather than just median splits. The analyses could then directly contrast and visualise participants who have scored the highest on a factor and those who have scored the lowest. This would enable us to characterize more targeted population groups.

Furthermore, as our participants belonged to a healthy group, another exciting research direction is to perform the same analyses for migraine/headache sufferers. This would offer the opportunity to relate the findings to previous research and to investigate whether the effects we have seen for a healthy group carry over to a clinically relevant group. In this way, these electrophysiological findings could potentially provide a biomarker to help in migraine diagnosis.

## 5. Conclusion

Ultimately, we have provided evidence that hyperexcitability in the visual cortex during the pattern-glare test exists in the healthy population and manifests as an increased overall amplitude for the clinically aggravating (Medium) stimulus. Sensitization to the clinically-relevant medium spatial-frequency stimulus was identified over a short time-frame, while habituation to the same stimulus was identified over a longer time-frame. Both this sensitization and habituation was dependent on susceptibility to headaches. These results suggest that this same experimental paradigm and analysis should be performed on a clinically diagnosed population.

## Data Availability Statement

The datasets used and/or analysed during the current study available from the corresponding author on reasonable request.

