## Appendix for "Headache-specific Hyperexcitation Sensitises and Habituates on different Time Scales: An Event Related Potential study of Pattern-Glare"


### Appendix 1: Ruling out eye confounds

To address the potential concern of eye movements driving the effects presented in this paper, analogues of each contrast were run on the eye electrodes to evaluate the possibility of any statistical contribution from eye confounds (Table A1.1). Table A1.2 lists the relevant electrodes and their location. Trials which were rejected during pre-processing for the non-eye related analysis were discarded from this analysis. This ensures a direct comparison between the eye electrodes and the set of trials used for analysis reported in this paper. Pre-processing and permutation test configurations applied in this analysis are the same as presented in the methods section.

MUA finds no clusters for all analyses on the eye electrodes (table A1.1).

| | **Partitions** | | **Onsets** | |
|---|---|---|---|---|
| **Contrast** | **Habituation** | **Sensitization** | **Habituation** | **Sensitization** |
| **Pure Time Effect** | All electrodes: None Found | All electrodes: None Found | All electrodes: None Found | All electrodes: None Found |

| | | | | |
|---|---|---|---|---|
| **Headache x Time (unorthogonalized)** | All electrodes: None Found | All electrodes: None Found | All electrodes: None Found | All electrodes: None Found |
| **Headache x Time (orthogonalized)** | All electrodes: None Found | All electrodes: None Found | All electrodes: None Found | All electrodes: None Found |

Table A1.1 Collective results for tests performed on the eye electrodes.

| **Electrode** | **Location** |
|---|---|
| EXG1 | 2cm above the right eye (aligned to centre of eye) |
| EXG2 | 2cm below the right eye (aligned to centre of eye) |
| EXG3 | 1cm right of the right eye (aligned to centre of eye) |
| EXG4 | 1cm left of the left eye (aligned to centre of eye) |
| EXG5 | On right mastoid behind right ear (reference) |
| EXG6 | On left mastoid behind left ear (reference) |
| HEOG | The difference of the horizontal electrodes |
| VEOG | The difference of the vertical electrodes |

*Table A1.2. Locations of eye and reference electrodes for the 128-channel BioSemi EEG system.*

**Appendix 2: Headache Questions**

For the assessment of headache symptoms, we selected relevant questions from a more general Headache and General Health questionnaire. Thus, we did not use the headache criteria specified by the International Headache Society (Arnold M., 2018) to diagnose migraine. These are criteria for clinical diagnosis and do not provide scale measures of headache proneness. However, the criteria rely heavily on headache intensity, duration, and frequency, and the presence of aura (covering a wide variety of sensory / motor disturbances), all of which were assessed by our questions.

The questions we used were worded as follows:

Headache Frequency: “Thinking about the number of headaches you have had over the past three months, how many headaches on average do you have per week/month/year? .........................................................”

Headache Duration: “How long does your typical headache last ..........................................................”

Headache Intensity: “On a scale of 1-10 (with 1 being the least), how bad was your worst headache? ............”

Aura: “Before or during your headache:

a. Do you ever feel nauseated or vomit? Yes [ ] No [ ]

b. Do you have a sensitivity to light? Yes [ ] No [ ]

c. Do you have a sensitivity to noise? Yes [ ] No [ ]

d. Does your eye become red or start watering or swell up on the painful side? Yes [ ] No [ ]

e. Does your eyelid close on the painful side? Yes [ ] No [ ]

f. Does your nose block up or run on the painful side? Yes [ ] No [ ]

g. Do you have motor weakness? Yes [ ] No [ ]

h. Do you feel pins and needles? Yes [ ] No [ ]

i. Do you have speech disturbances? Yes [ ] No [ ]

j. Is your hearing affected? Yes [ ] No [ ]

k. Is your vision affected? Yes [ ] No [ ]”

Participants had some freedom to specify headache frequency in terms of weeks, months, or years but their responses were converted to headaches per-year. For example, the response ‘2 per month’ was coded as 24. Similarly, participants specified headache duration in minutes, hours, or in some cases days, but duration was always coded in hours. Aura was coded as the total number of ‘yes’ responses to sub-questions a-k.

**Appendix 3: Factor Analysis**

Factor analysis seeks to derive (a small number of) uncorrelated factors from a collection of variables. Thus, the correlations between our factors were zero for the 39 participants who contributed viable questionnaire data. Rotation helps to clarify which variables load onto (belong to) each factor, as seen in the Rotated Component Matrix (Table A3.1) and helps with factor naming. Our factor scores were calculated in SPSS (IBM, NY) using the regression method in which all variables contribute to all factors according to the Component Score Coefficient Matrix (Table A3.2). For the headache and discomfort factors, the contribution from variables not obviously associated with these terms was minimal. The headache variables did contribute to the visual stress factor but were pitted against each other such that intensity and frequency contributed positively onto this factor, while duration contributed negatively. Since the headache variables correlate relatively strongly, the effect of these positive and negative weightings will be to cancel out the overall effect of headache on the VS factor. Although correlations between factors were zero by design, the correlation between discomfort ratings for the medium stimuli alone and headache frequency was 0.3 ($p$=.068) which, while not significant in our sample, is similar to previous measures.

**Table A3.1. Rotated Component Matrix**

| Rotated Component Matrix | | | |
|---|---|---|---|
| | Component | | |
| | 1 | 2 | 3 |
| VSQ | 0.636 | 0.035 | 0.062 |
| CHi | 0.845 | 0.212 | -0.074 |
| aura | 0.789 | 0.122 | 0.292 |
| H-duration | -0.470 | 0.764 | 0.040 |
| H-intensity | 0.366 | 0.716 | -0.116 |
| H-frequency | 0.305 | 0.795 | 0.041 |
| Discomfort | 0.119 | -0.028 | 0.977 |

**Table A3.2. Component Score Coefficient Matrix**

| | Component Score Coefficient Matrix | | |
|---|---|---|---|
| | | Component | |
| | 1 | 2 | 3 |
| VSQ | 0.303 | -0.055 | -0.021 |
| CHi | 0.401 | 0.018 | -0.172 |
| aura | 0.337 | -0.012 | 0.187 |
| H-duration | -0.332 | 0.51 | 0.135 |
| H-intensity | 0.108 | 0.372 | -0.128 |
| H-frequency | 0.048 | 0.433 | 0.036 |
| Discomfort | -0.065 | 0.013 | 0.934 |

### Appendix 4: Justification of factor analysis decomposition

An important point to consider is the reliability of the factor analysis we have performed. This section provides evidence for this reliability.

Firstly, we note Kaiser's rule, which suggests having at least five times as many observations as you have variables, will ensure a reliable fit. In our case, we have seven variables, so we meet the threshold of 35 participants (Kaiser; 1974), since we have fitted to 39 participants. Although, there are conflicting opinions on this issue, e.g. Costello & Osborne (2005).

However, all such "rules of thumb" are limited in their applicability, since they do not reflect the characteristics of any particular data set. Indeed, a data set with less noise variability is going to give more reliable factor analyses than one with more.

#### Background to Analysis

In justifying the reliability of our factor analysis, our basic approach is to compare the decomposition into factors and the uncertainty re. this decomposition, with what we would observe if we performed the only real alternative approach to a data-driven decomposition, such as factor analysis. This alternative is what we call a *flat average*.

More specifically, in order to identify electrophysiological features that correlate with condition-relevant features (e.g. headache susceptibility), we need regressors that classify our participants according to such features. One could just include regressors that are single variables from those we have collected, e.g. Headache-intensity, but that would throw away a good deal of information collected. One would also expect that aggregating across multiple response variables, would provide a more robust measure. Additionally, one wants an aggregation procedure that provides orthogonal regressors, which when entered into a regression model explain non-overlapping variability in the data. Without orthogonality of regressors, interpretation of findings is challenging.

As previously discussed, to do this, one is really left with two options, especially if one wants to restrict to a linear association of (condition-relevant) variables to components:

1) flat averaging of relevant variables, based upon an intuition of the condition-relevant features of each variable; and
2) data-driven selection of weighting coefficients by a procedure such as PCA or factor analysis.

We contend that there are only really two plausible flat average patterns as depicted in Figure A4.1. These patterns differ only in the assignment of aura to the visual stress or headache factors respectively. Phenomenologically and perhaps, mechanistically, one might consider aura to fit with visual stress, where susceptibility to spurious visual percepts and consequences of hyper-excitation load. Intuitively, one might also think that aura would associate with the headache variables (H-duration, H-intensity, H-frequency), since it is an experience that occurs before and during migraine headaches. Effectively, our data-driven approach has provided a clear answer to this question; see subsection "Comparison to flat average".

If one believed that our factor analysis was unreliable, it would generate a pattern of factor loadings that are substantially subject to noise. There are two obvious ways in which this noise might manifest:

1) It would cause the pattern of factor loadings to be quite different from the (a priori) intuitive flat average loadings.
2) It would lead to high variability/ uncertainty in the loadings that would be produced.

In the next two subsections, we explore whether there is any evidence of these manifestations of noise.

**Version 1 (aura in visual stress)**

| | Visual stress | Headache | Discomfort |
|---|---|---|---|
| VDS | 1 | 0 | 0 |
| CHi | 1 | 0 | 0 |
| Aura | 1 | 0 | 0 |
| H-duration | 0 | 1 | 0 |
| H-intensity | 0 | 1 | 0 |
| H-frequency | 0 | 1 | 0 |
| Discomfort | 0 | 0 | 1 |

**Version 2 (aura in headache)**

| | Visual stress | Headache | Discomfort |
|---|---|---|---|
| VDS | 1 | 0 | 0 |
| CHi | 1 | 0 | 0 |
| Aura | 0 | 1 | 0 |
| H-duration | 0 | 1 | 0 |
| H-intensity | 0 | 1 | 0 |
| H-frequency | 0 | 1 | 0 |
| Discomfort | 0 | 0 | 1 |

Figure A4.1: The two (intuitively) plausible flat-average associations of (condition-related) variables to components. CHi is the cortical hyperexcitability index; VDS is the Visual Discomfort Scale; H-duration is Headache duration and similarly for H-intensity and H-frequency. These two versions agree on the assignment of six out of seven (condition-related) variables, i.e. VDS and CHi to visual stress; H-duration, H-intensity and H-frequency to Headache; and Discomfort to Discomfort. The assignment of Aura (see rows in red) is the point of contention: it could plausibly be associated with either visual stress or headache, because it exhibits symptoms that one might associate with the CHi and VDS questionnaires, but occurs with headaches. An objective of our factor analysis was to provide a data-driven answer to the question of which component Aura should associate with.

**Comparison to flat average**

Our application of factor analysis gave the weighting coefficients across our seven variables in Figure A4.2.

Rotated Component Matrix

| | Component | | |
|---|---|---|---|
| | 1 | 2 | 3 |
| VDS | 0.636 | 0.035 | 0.062 |
| CHi | 0.845 | 0.212 | -0.074 |
| aura | 0.789 | 0.122 | 0.292 |
| H-duration | -0.470 | 0.764 | 0.040 |
| H-intensity | 0.366 | 0.716 | -0.116 |
| H-frequency | 0.305 | 0.795 | 0.041 |
| Discomfort | 0.119 | -0.028 | 0.977 |

Figure A4.2: factor loadings obtained through factor analysis on observed data.

importantly, these results are not substantially different to those we would obtain from flat averaging (see red rectangles). Although, as previously discussed, there is one condition-related variable (aura), for which there is uncertainty about its association (from an intuitive perspective) with components (see blue and green boxes).

Thus, our data-driven approach has provided loadings, which would correspond to the weighting matrix shown as version one (aura in visual stress), in figure A4.1.

The correspondence to our derived weightings is especially the case if one notes that the positive and negative headache variables on the first factor/component cancel in our factor loadings. That is, the overall effect of the three headache variables on the first factor is going to be much closer to zero than might be suggested by the absolute values of each of the three coefficients. This is because one is negative and the other two are positive.

In fact, it should be reassuring that a data driven approach like factor analysis, identifies a factor structure so close to one's intuitive expectations.

Effectively, the specific contribution of our data-driven approach is to provide a clear answer (loading in blue box is much higher than in green box) to the point of uncertainty we were faced with: which component should aura be associated with?

Aura correlating with visual stress is quite plausible with our population, who are sub-clinical. That is, it is likely that we have many participants who have headaches that are *neither* visually-induced nor associated with visual aura. For these participants, aura may be irrelevant to the frequency and features of their headaches. However, for others, the presence or absence of aura may be strongly correlated with the experience of visual stress-related percepts.

Thus, we would argue that the factor loadings we obtain are not speculative: the two possible approaches, data-independent or data-driven, are in the main very consistent in their outcomes, with the data-driven approach enabling us to determine the most appropriate component that aura should be allocated to in our participants.

**Variability investigation**

As previously discussed, if our factor analysis was especially subject to noise, we would expect this to manifest as instability in the factor loadings generated across replications. We seek to quantify this instability in this subsection and compare it to the instability found in the flat average approach, and we include a PCA decomposition for comparison.

*Methods*

To explore this instability, we bootstrapped our data, as one would do when quantifying error using confidence intervals. Thus, we started with our data represented as a 39 rows (participants) by 7 columns (variables) data structure. We then sampled each column with replacement to generate 50,000 (bootstrapped) surrogate data sets. On each such surrogate data set we ran three analyses, flat average, PCA and factor analysis, each of which projected the (39 x 7) surrogate data onto three components, giving a 39 x 3 projected surrogate data. More specifically, we performed the following procedures on each surrogate data:

1) *Flat average*: we projected onto three components, by applying the version 1 (flat average) weight matrix to the surrogate data, i.e. row *i* in the first column of the projected surrogate data was the average of the first three columns (VDS, Chi and Aura) of the *i*th row of the surrogate data, and so on for the next two columns. This gave us the *flat average projected data*.

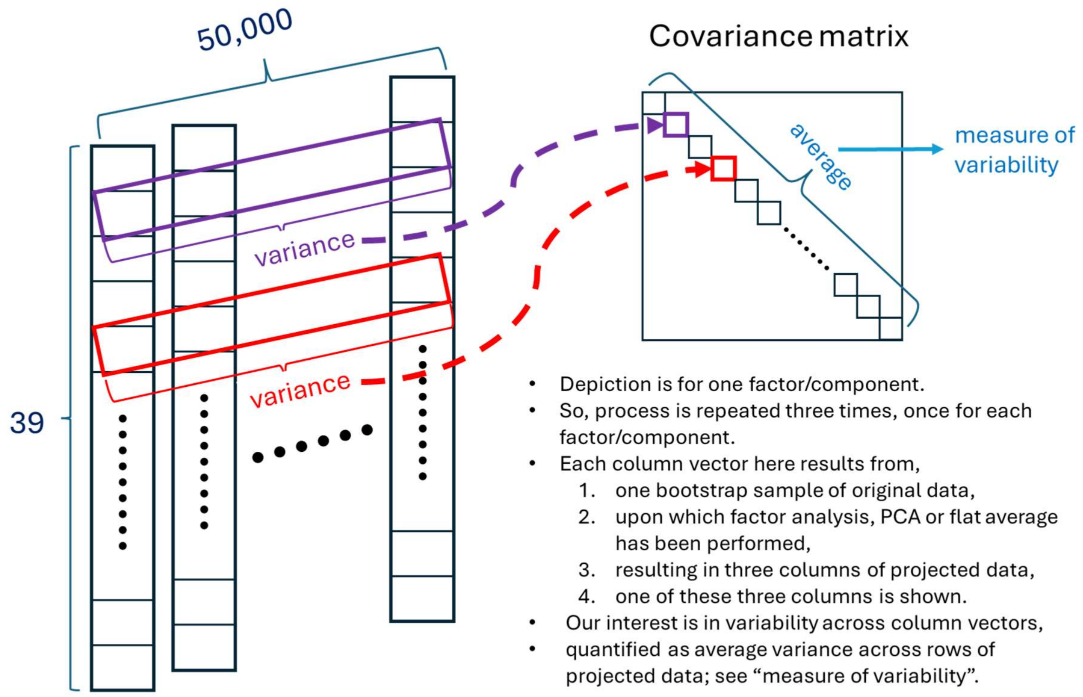


Figure A4.3: method for calculating variance induced by a particular method on projected data. 39 is the number of participants, 50,000 is the number of bootstrap resamplings. The main diagonal of the covariance matrix contains the variances of each row of 39x50,000 data structure. Each row corresponds to a (surrogate/bootstrapped) participant. The average of the variances across these rows (i.e. the trace of the covariance matrix divided by the number of rows) is a measure of variability/uncertainty resulting from projecting our data through either factor analysis, PCA or flat average loading/projection matrix.

2) *PCA*: we performed PCA decomposition on the surrogate data set, giving us a 7x3 matrix of component loadings/ weights. The surrogate data was then projected onto the three component dimensions using this matrix, giving us a 39x3 data set. This is the *PCA projected data*.
3) *Factor analysis*: we performed an exploratory factor analysis on the surrogate data set, giving us a 7x3 matrix of factor loadings/ weights. The surrogate data was then projected onto the three factor dimensions using this matrix, giving us a 39x3 data set. This is the *factor analysis projected data*.

So, for each of these procedures, we obtained 50,000 39 x 3 projected surrogate data sets. We are interested to quantify the variability across these 50,000. We do this separately for each of the three dimensions/components/factors projected onto, i.e. on 50,000 39 x 1 data structures. Thus, we are interested in the variability across these 50,000 column vectors, i.e. a 39 x 50,000 data structure; see figure A4.3. We quantify this variability by constructing the 39 x 39 covariance matrix across this 39 x 50,000 matrix; see figure A4.3. The trace of this covariance matrix is the sum of the variability across each of the 39 rows. Note, off-diagonal elements in the covariance matrix are close to zero, since the 50,000 is big (because the bootstrap sampling is random). Consequently, there is effectively no covariance across rows in figure A4.3, to be concerned about.

This gives us a sum of variances for each of the three components/ factors, which we turn into an average. If these three averages of variances are substantially bigger for factor analysis than it is for the flat average, it would suggest that the factor analysis adds uncertainty/error.

*Results*

In this section, we present the average along the trace from each covariance matrix across all bootstrapped samples for each of the three procedures for projecting onto three components. These results are presented in table A4.4.

| # Bootstraps (50,000) | 1 | 2 | 3 | Total Average Trace |
|---|---|---|---|---|
| **Averaging** | 0.6 | 0.91 | 0.88 | **2.39** |
| **PCA** | 2.62 | 1.57 | 0.95 | **5.14** |
| **EFA** | 0.91 | 0.91 | 0.89 | **2.71** |

Table A4.4: Average along trace from each covariance matrix across all bootstrap samples for each of the three procedures for projecting onto three components/factors (indicated 1, 2 and 3 here): (flat) Averaging, principal component analysis (PCA) and exploratory factor analysis (EFA). Additionally, the total averaged trace across all three dimensions is presented. These dimensions would be called components for PCA and factors for EFA.

As expected, the variability reduces across the PCA components – mathematically, it is placing each component at the highest variance dimension of the data remaining after projecting out previous components. Additionally, although to a lesser extent, variability reduces across factors, since more explanatory variables are again earlier. The total of the three average variabilities along the trace (the last column) is smallest for flat averaging, slightly higher for EFA and then substantially higher for PCA. This total is an estimate of the error/uncertainty associated with each procedure, and importantly, based on this analysis, experimental factor analysis does not substantially increase the error over that observed from our baseline procedure, (flat) averaging. That is, the flat average is the plausible alternative to using factor analysis and one might believe that its simplicity (and the fact that loadings are fixed) would make it substantially less susceptible to error. However, this is not what we see – with our bootstrapping procedure, factor analysis exhibited only a little more error variance than flat averaging. This increase in variability almost certainly arises because factors are ordered by their explanatory value, which relates to variance explained. We are only looking at the first three factors, which will necessarily have higher explained variance.

However, our central point is clear, even though the factor analysis loading matrix is derived from the data and changes between data sets, it does not add substantially more variability to the flat average, which is a fixed loading matrix. This suggests that, with our data, factor analysis is not a great source of added uncertainty.

*Algorithm*

To estimate variability in these methods we employed a bootstrapping procedure, i.e. using sampling with replacement. The algorithm is listed below:

1. **Initialisation:**
    a. Set the number of bootstrapped datasets to generate k=50,000.
    b. Define the number of factors/components for EFA/PCA (3).
    c. Instantiate three vectors of size (number_of_bootstraps, number_of_participants, 3) to store results for the averaging, PCA and EFA procedures.
2. **Bootstrapping Procedure:**
    a. For i from 1 to k:
        i. Generate a bootstrapped sample of size 39x7 by sampling with replacement from the original dataset of 39 participants and 7 variables.
        ii. For each method (averaging, PCA, EFA):
            1. Average
                a. Calculate the following three factors:
                    i. Visual Stress: Mean of VDS(z), Total Chi(z), and Aura(z)
                    ii. Headache: Mean of Headache Duration(z), Headache Intensity(z), and Headache Frequency(z)
                    iii. Discomfort: Discomfort Index(z)
                b. Store each of the three factors in the averaging vector i.e.
                    i. averaging_results(i, :, 0) = Visual Stress
                    ii. averaging_results(i, :, 1) = Headache
                    iii. averaging_results(i, :, 2) = Discomfort
            2. PCA:
                a. Apply Principal Component Analysis (PCA) to the bootstrapped dataset of size 39x7.
                b. Transform the dataset to a reduced dimensionality of size 39x3 using the first 3 principal components.
                c. Store each of the three components in the PCA vector i.e.
                    i. pca_results(i, :, 0) = PCA1
                    ii. pca_results(i, :, 1) = PCA2
                    iii. pca_results(i, :, 2) = PCA3
            3. EFA:
                a. Apply Exploratory Factor Analysis (EFA) to the bootstrapped dataset of size 39x7.
                b. Extract 3 factors and transform the dataset to a reduced dimensionality of size 39x3.
                c. Store each of the three components in the EFA vector i.e.
                    i. efa_results(i, :, 0) = Factor 1

ii. efa_results(i, :, 1) = Factor 2
iii. efa_results(i, :, 2) = Factor 3

3. **Compute the covariance matrix of each one of the methods**
    i. For each method, calculate the covariance matrix for its own respective factor, component, score i.e.
        1. for Visual Stress: covariance_matrix(averaging_results(:,:,0))
        2. for Headache: covariance_matrix(averaging_results(:,:,1))
        3. for Discomfort: covariance_matrix(averaging_results(:,:,2))
4. **Given the covariance matrix compute the average trace and present the results.**

### Appendix 5: Effects close to statistical significance

#### Orthogonalized Headache x Habituation for Partitions

MUA found no statistically significant clusters for the orthogonalized Headache x Habituation for Partitions effect, the regressor for which is shown in Figure A5.1.1. Non-significant results are reported in table A5.1.1.

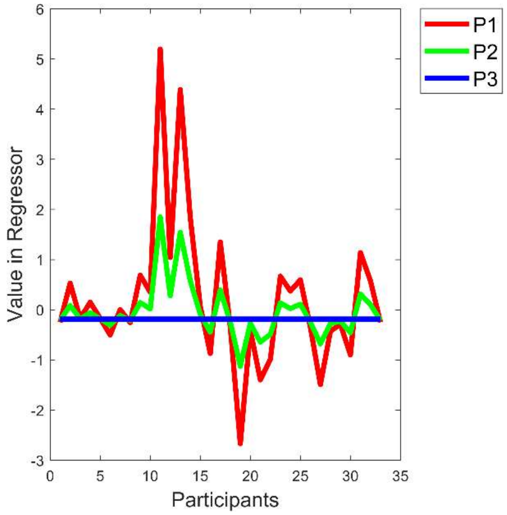


*Figure A5.1.1: Orthogonalized Headache x Habituation for Partitions regressor. The x-axis represents the participants and the y-axis the design matrix value assigned to them for each Onset.*

Figure A5.1.2*(c)* displays the grand average time-series for the most continuously significant electrode in the cluster, divided into high and low according to a median split on participant scores on the orthogonalized Headache factor. The grand average time-series on the first and second rows of Figure A5.1.2*(c)* show no clear habituation during the window (see two vertical lines) of the first (most significant) cluster (blue in panel a), however a difference in amplitudes is observed (higher for the high group, lower for the low group). Therefore, this suggests that the effect is driven mostly by hyperexcitation in the high group when compared to the low group, and not necessarily habituation through the partitions.

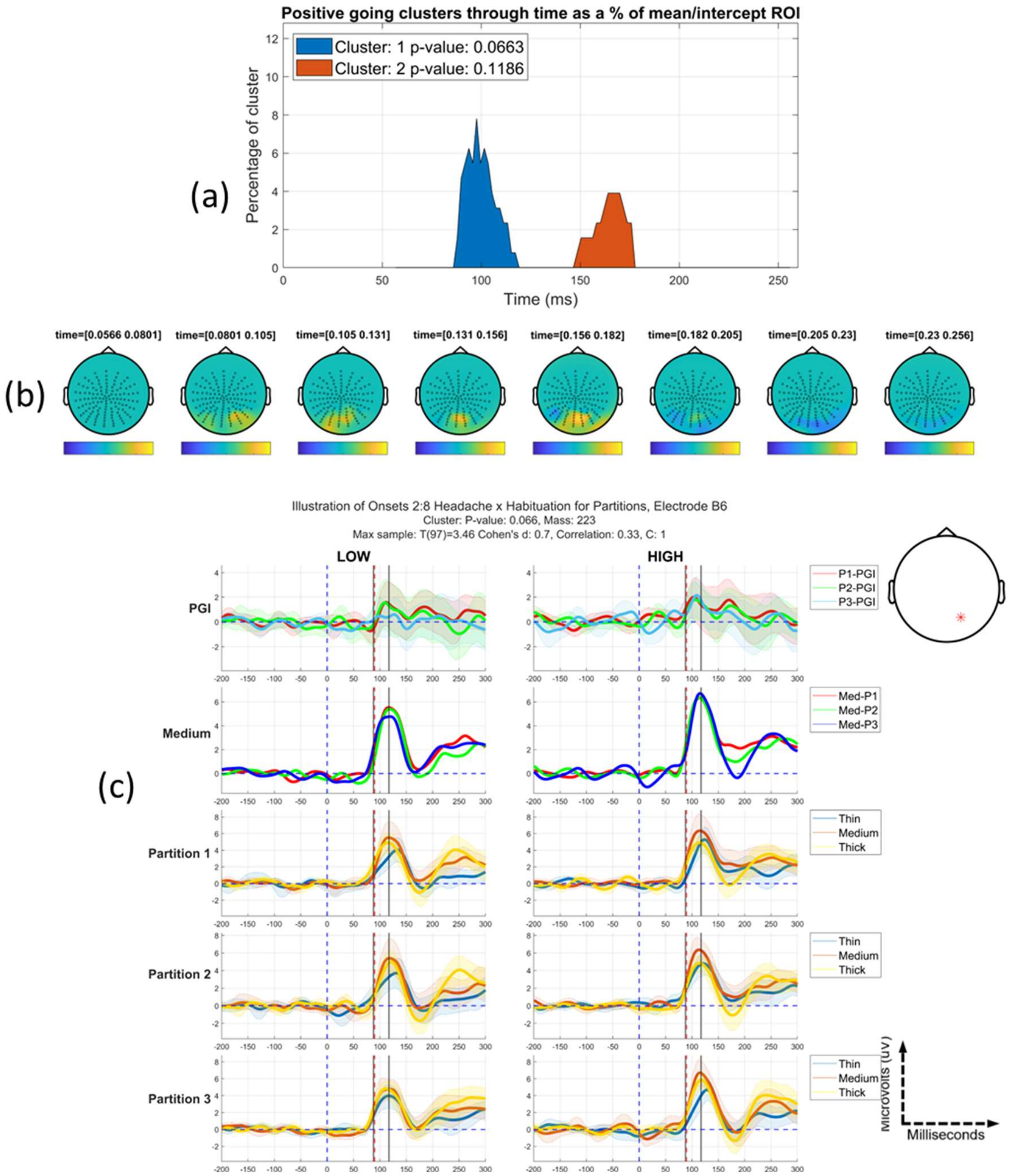


*Figure A5.1.2 Orthogonalized Headache x Habituation for Partitions* effect: *(a) Positive going clusters through time as a proportion of the mean/intercept region-of-interest. (b) Topographic maps through bounding time window, 56-256ms after stimulus onset, with time step between each map approximately 25ms. Positive-going clusters are observed at posterior electrodes, specifically in the region of 95ms-175ms. (c) Grand average time-series of the PGI and individual conditions at the electrode that is the most continuously significant through time for the most significant cluster (blue in panel (a)). A median split was taken and plots on the left side are for those low on the headache factor and the right side for those high on the factor. The shaded regions represent 95% confidence intervals for each time-series. The black lines indicate the start and end of the most significant cluster through time, with the red dashed line indicating the peak of the effect. Partitions are indicated by P1, P2 and P3. Med_Pi indicates the medium stimulus for partition i. C: j indicates the cluster number.*

| *Orthogonalized Headache x Sensitization for Onsets* | | |
|---|---|---|
| *<u>Statistic</u>* | *<u>PC1</u>* | *<u>PC2</u>* |
| *p* | 0.0663 | 0.1186 |
| *Most sustained electrode* | B6 | A28 |
| *Time of peak of most sustained* | 90ms | 168ms |
| *Max T(34)-value* | 3.5 | 2.6 |
| *Cluster Stat* | 223 | 117 |

*Table A5.1.1 Collective results for positive (+1) tail of the orthogonalized Headache x Habituation for Partitions effect. For each effect that crossed the cluster forming threshold, the following values are presented: the significance probability (p-value, FWE-corrected); the electrode where the effect is most sustained through time; the time of the peak for the most sustained electrode; the t-value at that peak; and the summed t-values of the significant cluster. Clusters are ordered (from left to right) from most to least significant. PC = positive cluster found within the ROI. * indicates p≤0.05.*

### <u>Orthogonalized Headache x Habituation for Partitions x Sensitization for Onsets</u>

MUA found no statistically-significant clusters for the Orthogonalized Headache x Habituation for Partitions x Sensitization for Onsets effect. The regressor for this interaction is presented in Figure A5.2.1 and results are summarised in table *A5.2.1* The largest effect is visualised in figure *A5.2.2*. The most direct way to understand the effect observed, is to concentrate on the PGI rows in panel (c): the first (partition 1), third (partition 2) and fifth (partition 3) rows (panels with grey backgrounds). Although no cluster is significant, in the time-period of the cluster closest to significant, there is some evidence of partitions differentially (across high and low groups) modulating the effect on onsets. For example, we do see a sensitization effect across onsets in the top right plot in panel (c) (i.e., High group, partition 1), which is not observed for any other partition in the High or Low group. However, the effect is far from significant.

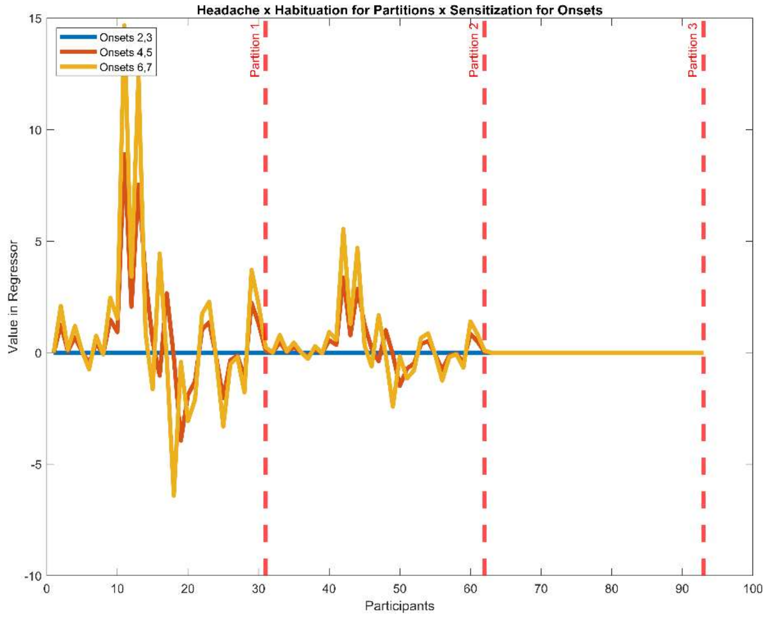


*Figure A5.2.1:* Orthogonalized Headache x Habituation (for Partitions) x Sensitization (for Onsets) regressor: *the x-axis represents participants, all of which appear three times, once for each partition. The y-axis gives the design matrix value assigned to each participant for each onset in each partition. Onsets are indicated by the blue, orange, and yellow lines (2:3; 4:5; 6:7) with partitions segmented by the red dashed lines. Note, in the final partition (number 3), the red and blue lines are "under" the yellow line.*

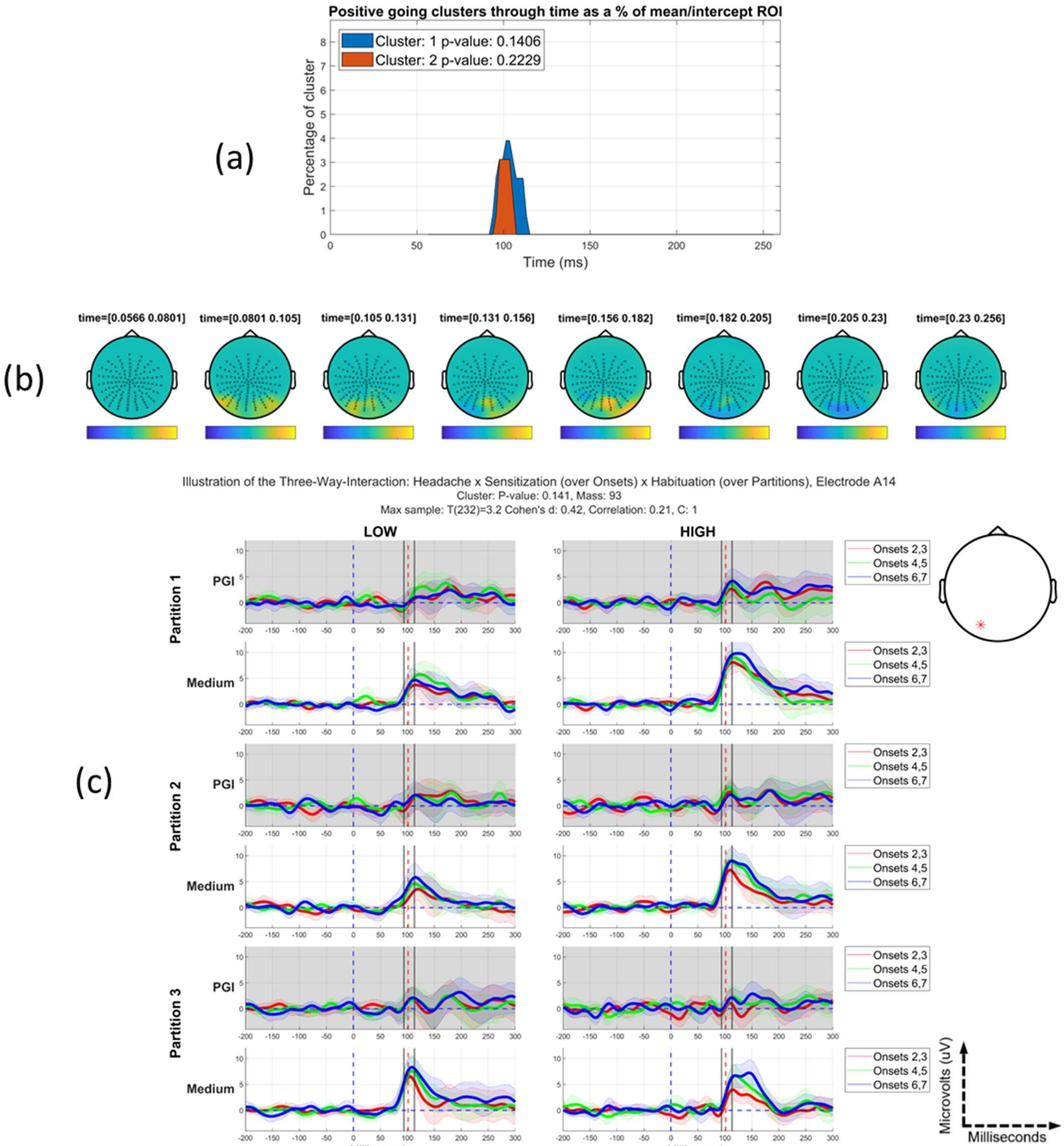


*Figure A5.2.2* (Orthogonalized) Headache *x Habituation (for Partitions) x Sensitization (for Onsets)* effect: *(a) Positive going clusters through time as a proportion of the mean/intercept region-of-interest. (b) Topographic maps through bounding time window, 56-256ms after stimulus onset, with time step between each map approximately 25ms. The cluster with the lowest p-value is positive-going at left posterior electrodes in the region of 85-110ms. (c) Grand average time-series of the PGI and Medium stimulus at the electrode that has the lowest accumulated p-value through time in the cluster closest to significance. The shaded regions represent 95% confidence intervals for each time-series. A median split was taken and plots on the left side are for those low on the headache factor and the right side for those high*

*on the factor. The black lines indicate the start and end of the cluster closest to significance with the red dashed line indicating the peak of the effect. Partition 1 is indicated by the first two rows, partition 2 the second two and the last two partition 3. Within each partition, the PGI and medium stimulus is highlighted for onsets 2:3, 4:5 and 6:7.*

| *Orthogonalized*<br><br>*Headache x Habituation for Partitions x Sensitization for Onsets* | | |
|---|---|---|
| ***Statistic*** | ***PC1*** | ***PC2*** |
| *p* | 0.1406 | 0.2229 |
| *Most sustained electrode* | A14 | A30 |
| *Time of peak of most sustained* | 101ms | 152ms |
| *Max T(34)-value* | 3.2 | 2.5 |
| *Cluster Stat* | 93 | 73 |

*Table A5.2.1 Collective results for the positive (+1) tail of the orthogonalized Headache x Habituation for Partitions x Sensitization for Onsets interaction. For each effect that crossed the cluster forming threshold, the following values are presented: the significance probability (p-value, FWE-corrected); the electrode where the effect is most sustained through time; the time of the peak for the most sustained electrode; the t-value at that peak; and the summed t-values of the significant cluster. Clusters are ordered (from left to right) from lowest to highest p-value; PC = positive cluster found within the ROI. * Indicates p≤0.05.*
